# TogoMCP: Natural Language Querying of Life-Science Knowledge Graphs via Schema-Guided LLMs and the Model Context Protocol

**DOI:** 10.64898/2026.03.19.713030

**Authors:** Akira R. Kinjo, Yasunori Yamamoto, Samuel Bustamante-Larriet, Jose-Emilio Labra-Gayo, Takatomo Fujisawa

**Author notes:** Corresponding author. (Y.Y.). These authors contributed equally to this work (Akira R. Kinjo, Yasunori Yamamoto).

## Abstract

Querying the RDF Portal knowledge graph maintained by DBCLS—which aggregates approximately 60 life-science databases—requires proficiency in both SPARQL and database-specific RDF schemas, placing this resource beyond the reach of most researchers. Large Language Models (LLMs) can, in principle, translate natural-language questions into executable SPARQL, but without schema-level context, they frequently fabricate non-existent predicates or fail to resolve entity names to database-specific identifiers. We present TogoMCP, a system that recasts the LLM as a protocol-driven inference engine orchestrating specialized tools via the Model Context Protocol (MCP). Two mechanisms are essential to its design: (i) the MIE (Metadata-Interoperability-Exchange) file, a concise YAML document that dynamically supplies the LLM with each target database’s structural and semantic context at query time; and (ii) a two-stage workflow separating entity resolution via external REST APIs from schema-guided SPARQL generation. On a benchmark of 50 biologically grounded questions spanning five types and 23 databases, TogoMCP achieved a large improvement over an unaided baseline (Cohen’s *d* = 1.82, Wilcoxon *p* < 0.001), with win rates exceeding 80% for question types with precise, verifiable answers. An ablation study shows that all component configurations deliver significant improvements, with MIE schema files providing the largest marginal contribution on mean per-question score (Δ = +0.50 relative to a no-MIE condition, two-sided Wilcoxon *p* = 0.067; 90% bootstrap CI [+0.04, +0.94] excludes zero); a one-line instruction to load the relevant MIE file recovers the same mean improvement as a full procedural protocol, while the protocol additionally reduces downside risk (loss rate 1.6% vs. 4.8%, Fisher *p* = 0.036). These results suggest a general design principle: concise, dynamically delivered schema context is more valuable than complex orchestration logic for mean-score performance, while procedural guidance plays a complementary role in narrowing variance.

**Database URL:** https://togomcp.rdfportal.org/

## Introduction

The life sciences depend on a growing ecosystem of specialized databases covering nucleotides, genes, proteins, chemical compounds, genetic variants, and more. Because these data are large, heterogeneous, and continuously updated, many providers have adopted the Resource Description Framework (RDF) (1) as a common representation since the 2000s; UniProt (2) alone now exceeds 217 billion triples. The RDF Portal (3) maintained by the Database Center for Life Science (DBCLS) aggregates approximately 60 such datasets—spanning genomics, proteomics, molecular biology, and biomedicine (UniProt (2), ChEMBL (4), MeSH (5), GO (6), among others)—into an interlinked knowledge graph queryable via SPARQL (7). In principle, SPARQL enables precise, cross-database retrieval; in practice, the steep learning curve of both the query language and the database-specific RDF schemas places this resource beyond the reach of most life science researchers. Access logs for the DBCLS RDF Portal (April 2026) record over 10 million queries per month across six endpoints; an estimated 0.4% are human-initiated (DBCLS, personal communication), with the remainder generated by automated scripts and programmatic pipelines. Within that small fraction of human-initiated requests, only a subset is likely to represent users authoring SPARQL queries directly— the remainder being routed through interactive query browsers, prepared example queries, or other forms of human-mediated access that do not require SPARQL fluency. The aggregate picture is that direct SPARQL access remains far from a broadly used interface for the life-science research community.

Large Language Models (LLMs) (8) are a natural candidate for bridging this gap: their language understanding and general reasoning capabilities could translate a researcher’s question directly into executable SPARQL (9; 10). In our initial experiments, however, LLMs operating without an external context frequently produced syntactically valid but semantically incorrect queries—fabricating non-existent predicates, misusing database-specific vocabulary, or failing to resolve natural-language entity names (e.g., “Aspirin”) to the database-specific identifiers required by the knowledge graph (e.g., PubChem Compound ID 2244). The root cause is not a deficit of reasoning ability but a lack of access to up-to-date, schema-level knowledge that was never part of the model’s pre-training data.

Recent work has begun to address this challenge along two main axes. One line of research fine-tunes open LLMs on domain-specific question–SPARQL pairs: Rangel et al. (11) fine-tuned OpenLLaMA on the Bgee gene-expression knowledge graph, demonstrating that semantic clues embedded in SPARQL (meaningful variable names, inline comments) can improve generation accuracy by up to 33%, but the approach requires curating training data for each target database. A second line leverages retrieval-augmented generation (RAG) to supply schema context at query time without retraining. Emonet et al. (12) built a RAG pipeline over SIB bioinformatics endpoints that retrieves ShEx (Shape Expressions) class schemas and example queries via embedding similarity, with iterative validation against VoID descriptions; their extended system, SPARQL-LLM (13), adds systematic evaluation on both multilingual benchmarks and real-world federated knowledge graphs. Doulaverakis et al. (14) applied a similar prompting strategy to medical Linked Open Data, and Ali et al. (15) showed that grounding LLM outputs in an ontology-based RDF knowledge graph eliminates hallucinations that standalone models produce in clinical question answering. These systems, however, share two limitations: they target a single endpoint ecosystem or a small set of homogeneous databases, and they conflate entity resolution with query generation into a single prompting step.

**TogoMCP** addresses both limitations by recasting the LLM as a protocol-driven inference engine that orchestrates specialized external tools via the Model Context Protocol (MCP)^1^, a standardized framework developed by Anthropic for connecting LLMs to external resources, including Web service APIs and local data schemas (9). To cover the breadth of 28 heterogeneous databases, the system introduces the *MIE (Metadata-Interoperability-Exchange) file*, a concise YAML document that dynamically supplies the LLM with the structural and semantic context of each RDF database at the point of need. To decouple entity resolution from query generation, it adopts a two-stage hybrid workflow that separates *entity resolution* (rapid identifier lookup via specialized REST APIs) from *knowledge graph exploration* (LLM-generated SPARQL guided by the MIE file). This separation allows the LLM to focus its reasoning on graph traversal using concrete, verified identifiers rather than attempting brittle text-based entity matching within SPARQL itself.

Figure 1 illustrates how TogoMCP works. First, an agent accepts a question given by its user. Any implementation can serve as the agent, provided it supports MCP; here, we used Claude Sonnet 4.5 as a reference. The agent analyzes the natural-language question, extracts biologically relevant key terms, and selects the appropriate databases (Step 1). In the figure, the question is *“Show me any compounds binding to human TP53”*, and the database and the key term are *ChEMBL* and *TP53*, respectively. The key terms are searched across the relevant databases via their corresponding search APIs to retrieve entity IDs (Step 2). Once the IDs are obtained, the agent is expected to obtain the MIE files for the relevant databases (Step 3). In the Figure, the agent requests the ChEMBL MIE file. The next step is to build a SPARQL query to retrieve relevant data for the obtained ID by referencing the MIE (Step 4). Depending on the results returned by TogoMCP, the agent may reconstruct a query to refine the results and present them to the user.

**Fig. 1.**
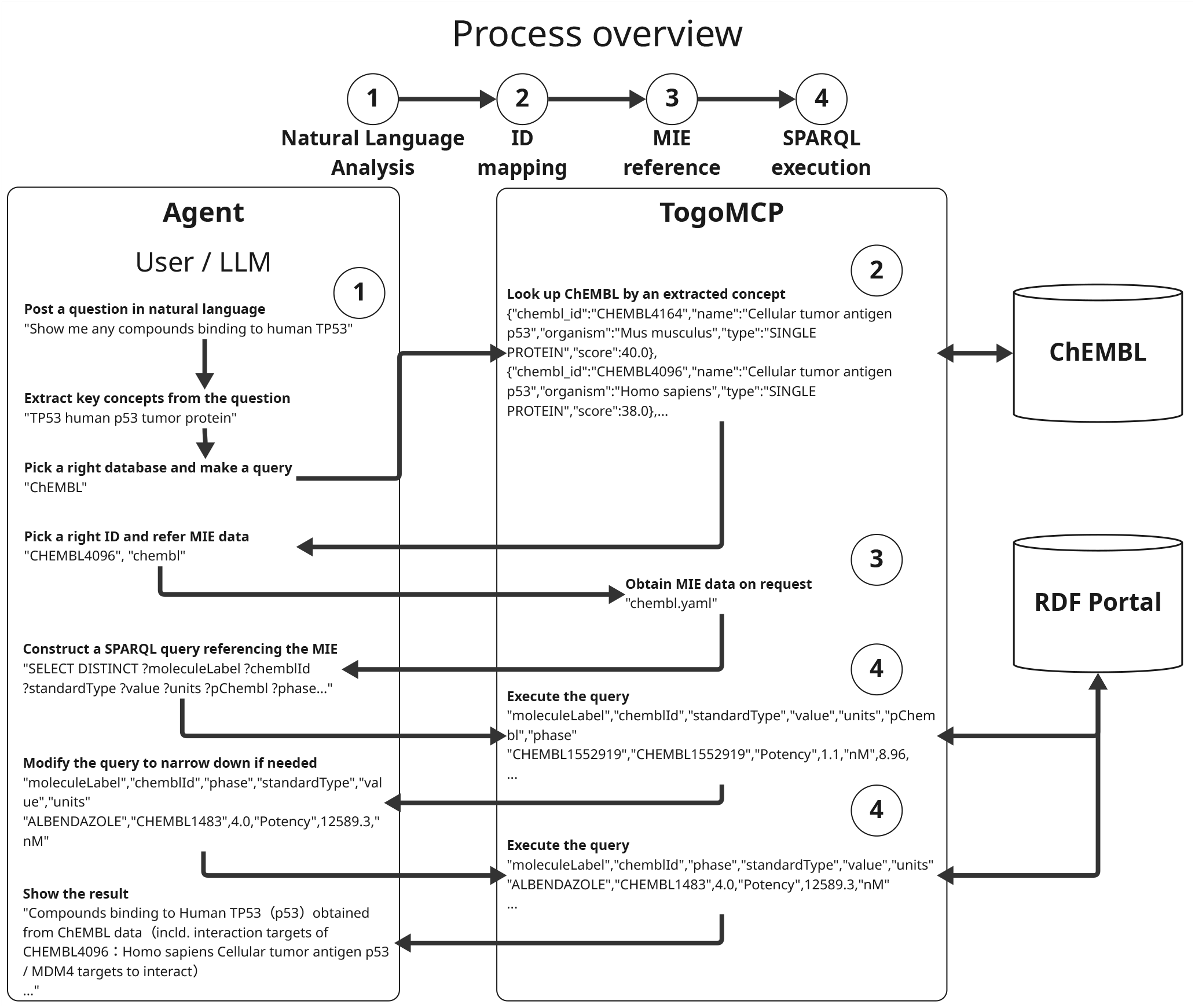
Overview of the TogoMCP workflow for the example query “Show me any compounds binding to human TP53.” The four stages proceed left to right: (1) natural language analysis to extract key concepts, (2) entity resolution via external search APIs to obtain database-specific identifiers, (3) retrieval of the target database’s MIE file for schema context, and (4) schema-guided SPARQL construction and execution against the RDF Portal. The left column traces the agent’s reasoning; the right column shows the corresponding TogoMCP tool calls and responses. Alt text: Flowchart showing the four-stage TogoMCP workflow in three columns: the agent (left) iteratively analyses the question, picks a database, requests an MIE file, and constructs SPARQL; TogoMCP (centre) returns ranked search hits, the MIE file, and SPARQL results; ChEMBL and the RDF Portal (right) are the external data sources.

The central contribution of TogoMCP is not the observation that schema knowledge is useful—that much is established by prior fine-tuning and RAG-based approaches (11; 13)—but the design of the *MIE file* as a compact, standardized, and database-portable unit of schema context, and its empirical characterization as the necessary and sufficient condition for LLM-generated SPARQL to succeed across 28 heterogeneous databases. The relevant empirical question is therefore not *whether* schema context helps, but *which form* is required and how much additional orchestration complexity is warranted. We evaluate TogoMCP on a benchmark of 50 biologically grounded questions spanning five question types and all 23 databases supported at the time of benchmark creation. The system achieves a large improvement over an unaided LLM baseline (Cohen’s *d* = 1.82), with gains concentrated in information recall and precision, and win rates exceeding 80% for question types with precise, verifiable answers. An ablation study shows that all component configurations deliver significant improvements, with MIE schema files providing the primary marginal contribution to the mean: removing them reduces the per-question improvement by 0.50 points (two-sided Wilcoxon *p* = 0.067; 90% bootstrap CI [+0.04, +0.94] excludes zero). A one-line system-prompt instruction to consult the relevant MIE file recovers the same mean-score improvement as an elaborate behavioural protocol; the protocol’s additional value is a complementary one, narrowing the variance of outcomes and reducing the per-evaluation loss rate roughly threefold (1.6% vs. 4.8%; Fisher *p* = 0.036). This result points to a general design principle for tool-augmented LLM systems: concise, dynamically delivered schema context is the highest-leverage component for mean performance, while procedural orchestration earns its place by managing the worst-case rather than by raising the average. The remainder of this paper details the architecture and methodology of TogoMCP, presents the benchmark evaluation and ablation results, and discusses implications for natural language interfaces to structured knowledge bases.

## Materials and Methods

### System Architecture and Environment

The system is implemented in Python (≥3.11) and built on the FastMCP library (version ≥3.0.0), which provides a standardized Model Context Protocol interface between the LLM and external resources. The LLMs used throughout are Anthropic’s Claude Sonnet 4.5 (generation) or Opus 4.7 (evaluation). The codebase comprises six modules organized into two functional layers:

1. **SPARQL layer**: Provides endpoint configuration, query execution (`run_sparql`), database discovery (`find_database_s`, `list databases`, `get_sparql_endpoints`), MIE file retrieval (`get_MIE_file`), and the TogoMCP Usage Guide (`TogoMCP_U_sage_Guide`).
2. **Web API layer**: Wraps external REST services for high-speed entity resolution and cross-database identifier mapping. Database-specific search tools cover UniProt, ChEMBL (targets, molecules), PubChem, PDB (via PDBj), MeSH, Reactome, and Rhea. NCBI E-utilities (16) support keyword search across eight NCBI databases (Gene, Taxonomy, ClinVar, MedGen, PubMed, PubChem Compound/Substance/BioAssay) with field-tag validation. TogoID (17) provides route discovery (`getAllRelatio n`), identifier conversion (`convertId`), and pre-conversion counting (`countId`).

An administrative module supports the generation of MIE files. The NCBI and TogoID sub-servers are mounted onto the primary FastMCP server at startup via the `mount()` mechanism, yielding a single MCP endpoint that exposes all tools to the LLM.

In addition to the TogoMCP server, the LLM client connects to three external MCP servers: **PubMed** (literature search, article metadata, and full-text retrieval), **OLS4** (18) (Ontology Lookup Service; term search, class hierarchy traversal via `g etAncestors`/`getDescendants`, and ontology browsing), and **PubDictionaries** (19) (controlled vocabulary and dictionary lookup). These complement TogoMCP’s tools by providing standardized access to biomedical literature, ontology term normalization (e.g., resolving keywords to GO or MeSH IRIs—Internationalized Resource Identifiers), and terminology services without requiring SPARQL. When these external services are unavailable, the Usage Guide specifies documented fallbacks: `search_mesh_descriptor` substitutes for OLS4 term lookup, and `ncbi_esearch` substitutes for PubMed.

Endpoint configuration is loaded from a CSV file, mapping each database to its SPARQL URL, endpoint group name, and preferred keyword-search tool. The system interfaces with 28 RDF databases (2; 4; 6; 5; 20; 21; 22; 16; 30; 31; 32; 33; 34) hosted across eight SPARQL endpoints (Table 1). Databases sharing an endpoint (23) can be queried with a single SPARQL statement using `GRAPH` clauses; databases on different endpoints require sequential queries bridged by TogoID (17) or NCBI cross-references. Of the 28 databases, 16 have dedicated REST entity-search tools (e.g., `search_uniprot_entity`, `search_chem_bl_target`, `ncbi_esearch`) that support fast, ranked identifier lookup before SPARQL is issued; the remaining 12 (marked† in Table 1) rely on SPARQL text search as a fallback, which the MIE anti-patterns section flags as a last resort due to its lower precision.

**Table 1.**
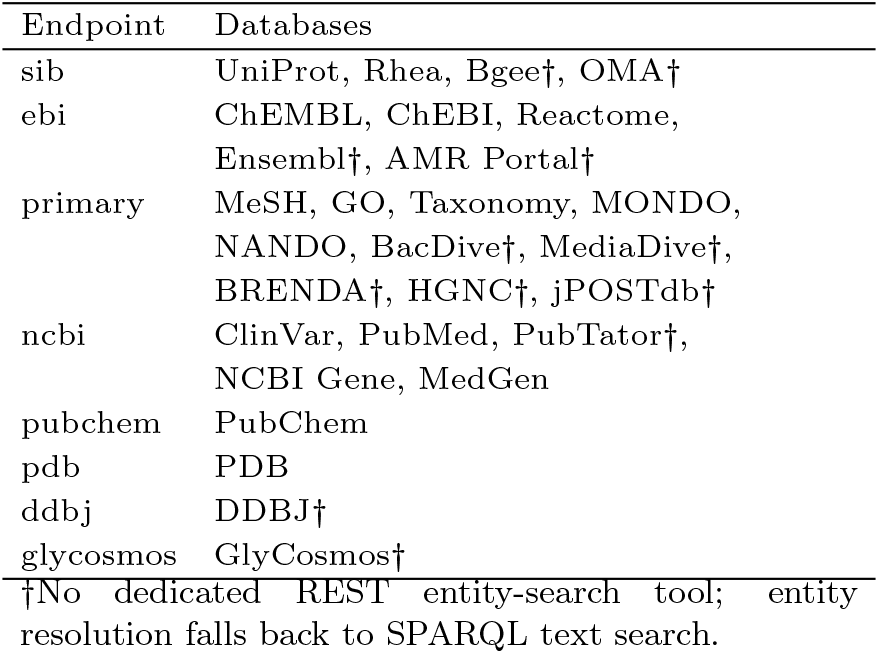
SPARQL endpoint groups and their constituent databases. All RDF Portal endpoints share the base URL https://rdfportal.org/{endpoint}/sparql, except GlyCosmos (29) (ts.glycosmos.org/sparql).

| Endpoint | Databases |
| --- | --- |
| sib | UniProt, Rhea, Bgee†, OMA† |
| ebi | ChEMBL, ChEBI, Reactome, Ensembl†, AMR Portal† |
| primary | MeSH, GO, Taxonomy, MONDO, NANDO, BacDive†, MediaDive†, BRENDA†, HGNC†, jPOSTdb† |
| ncbi | ClinVar, PubMed, PubTator†, NCBI Gene, MedGen |
| pubchem | PubChem |
| pdb | PDB |
| ddbj | DDBJ† |
| glycosmos | GlyCosmos† |

†No dedicated REST entity-search tool; entity resolution falls back to SPARQL text search.

Database selection was guided by the goal of covering the data types that life-science researchers most routinely query: molecular sequence and function (UniProt, NCBI Gene, Ensembl), chemical compounds and drug targets (ChEMBL, ChEBI, PubChem), metabolic reactions and pathways (Rhea, Reactome, BRENDA), disease and clinical variant information (MONDO, NANDO, ClinVar, MedGen, MeSH), taxonomic and microbial diversity (NCBI Taxonomy, BacDive, MediaDive), structural biology (PDB), glycomics (GlyCosmos), gene expression (Bgee), orthology (OMA), antimicrobial resistance surveillance (AMR Portal), nucleotide sequence archives (DDBJ), proteomics (jPOSTdb), and gene nomenclature (HGNC). An additional technical requirement is that each database exposes a stable, queryable SPARQL endpoint with sufficient schema documentation to construct a MIE file.

Beyond the choice of which tools to expose, several design decisions at the level of individual tool signatures and error behaviour reduce the rate at which the LLM stalls or wastes its tool-call budget on recoverable failures. Tool descriptions are written as imperative protocol contracts rather than as declarative documentation: parameter values that the server can enumerate (such as database and endpoint identifiers) are listed inline in the description, and descriptions of error-prone operations include explicit anti-retry guidance, so decision logic that would otherwise require the LLM to consult external documentation is folded into the tool surface itself. When a remote API call fails, the wrapping tool catches the transport-layer error and returns a structured *Error: …* string with in-band recovery guidance (for example, “retry once after a brief delay; if it keeps failing, fall back to SPARQL with the following pattern”), rather than raising an exception that would disrupt the MCP session.

The server also tolerates the parameter-name variations that LLMs naturally produce. Database-taking tools accept `database`, `db`, and `dbname` interchangeably, and `run sparql` accepts both `query` and `sparql_query`; a server-wide middleware silently strips unknown keyword arguments from any `search *` call before validation, absorbing the LLM’s tendency to invent filter parameters (e.g. `organism=`, `reviewed=`) that the actual tool signature does not declare. Discoverability is concentrated in two complementary tools: `find_databases` returns a relevance-ranked short list by matching the question against a curated `keywords` field maintained in each MIE file, so that the most relevant databases appear at the top regardless of catalog size; and `get_graph_list` filters out internal system graphs (Virtuoso/OpenLink bookkeeping graphs such as `virtrdf` and `ldp`) and ranks remaining data graphs by case-insensitive substring match against the queried database name, so that the named-graph URI required by the SPARQL `FROM` clause appears at the top of the response rather than being buried among co-tenant graphs on multi-database endpoints.

Onboarding a new database requires approximately 2–4 hours of expert effort, following the same four-phase pipeline described in section “Context Provisioning via MIE Files”: automated schema discovery, LLM-assisted MIE drafting, validation against the live endpoint, and expert review. Priority for future additions is informed by usage-log data from the DBCLS query platform, which reflects the actual database demand of the Japanese life-science research community. The MIE specification (v2.1) is endpoint-agnostic and applies to any SPARQL-accessible RDF database; the expansion from 23 databases at initial submission to 28 at revision demonstrates that the onboarding pipeline scales in practice without requiring changes to the core server.

### Context Provisioning via MIE Files

A recurring failure mode observed during early development was the LLM’s generation of syntactically valid but semantically incorrect SPARQL—for example, fabricating non-existent predicates or misusing database-specific vocabulary. To address this, we developed the **MIE (Metadata-Interoperability-Exchange)** file format: a YAML document that supplies the LLM with the structural and semantic context required to query a given RDF database correctly.

One MIE file was created for each of the 28 databases following a four-phase generation pipeline. **Phase 1 (Schema Discovery)** explores the endpoint’s schema through a prescribed series of SPARQL queries, identifying entity classes, IRI patterns, and representative examples. **Phase 2 (LLM-Assisted Drafting)** synthesises the discovery findings into all eleven required YAML sections, following a structured authoring protocol that prioritises structured lookups over text search and records silent-failure traps in `critical_warnings`. **Phase 3 (Validation)** verifies all SPARQL examples and RDF entries against the live endpoint, cross-checks data statistics, and applies section-level audits introduced in MIE specification v2.1. **Phase 4 (Expert Review)** provides a final domain check for biological accuracy before the file is committed.

Each MIE file contains the following sections:

1. `schema_info`: Database metadata: title, description, SPARQL endpoint URL, named graphs, available keyword-search tools, version, license, and backend type.
2. `shape_expressions`: A focused subset of Shape Expressions (ShEx) (24) defining the RDF structure of major entity classes and their properties, using the actual prefixes and IRIs of the database—the primary mechanism by which the LLM learns database-specific vocabulary such as `up:reviewed` (UniProt RDF vocabulary prefix `up:`) or `cco:standardType` (ChEMBL Core Ontology prefix `cco:`)^2^.
3. `sample_rdf_entries`: Three representative RDF triple samples illustrating typical data patterns.
4. `sparql_query_examples`: Seven tested SPARQL queries at three complexity levels—2 basic, 3 intermediate, 2 advanced—with at least two using specific IRIs or `VALUES` clauses and at most one using text search.
5. `cross_database_queries`: Two to three examples of `GRAPH`-based joins for databases sharing an endpoint.
6. `cross_references`: Documentation of cross-reference patterns to external databases.
7. `data_statistics`: Verified entity counts and coverage percentages, each with a verification date and method (e.g., direct COUNT query).
8. `anti_patterns` **/** `common_errors`: Three to four documented failure patterns with corrected alternatives, e.g., using text search when a structured predicate exists, or circular reasoning with search-result IDs.

The MIE file is delivered to the LLM on demand via the `get_MIE_file` tool. A typical file is 400–600 lines (700–900 for complex databases such as UniProt or ChEMBL), which is compact enough to fit within the LLM’s context window while providing sufficient detail for accurate query generation. The `list_databases` tool returns the `schema_info` section of all 28 files, enabling the LLM to select relevant databases before loading full MIE files.

### Protocol-Driven Control: The TogoMCP Usage Guide

The TogoMCP Usage Guide is an empirically devised execution protocol, delivered to the LLM as the first tool call of every session via the `TogoMCP_Usage_Guide` tool. It governs all subsequent tool selection, sequencing, and failure handling. The guide was iteratively refined using performance data from 150 evaluated question–answer pairs and encodes the following key strategies.

#### Question analysis before tool use

The LLM must classify the query into one of five types—verification, enumeration, comparative, synthesis, or exploration—before invoking any tool. For each type, the guide specifies a SPARQL budget (e.g., 1–2 calls for verification, 3–4 for comparative), a total tool-call budget (optimal range: 6–15), and a step-by-step workflow template.

#### Mandatory workflow

Every query follows a fixed sequence: (1) `find_databases` for database discovery (with `list_databases` as a full-catalog fallback); (2) entity resolution via specialized search tools or NCBI E-utilities; (3) `get_MIE_file` for each database to be queried; (4) targeted `run_sparql` calls; and (5) synthesis of the final answer. Cross-database bridging via TogoID must be planned within the first 3–5 tool calls when databases reside on different endpoints.

#### SPARQL discipline

Empirical analysis showed that questions with at most two consecutive SPARQL calls scored 17.81/20 on average, whereas those with three or four consecutive calls dropped to 16.55. The guide, therefore, enforces a hard limit of two consecutive SPARQL calls, after which the LLM must pivot to an alternative strategy: simplifying the query, switching to a search tool, or using TogoID to bridge to a different database. A total of 1–3 SPARQL calls per question is identified as the optimal range; beyond seven, scores decline to baseline-equivalent levels.

#### Tool prioritization

The guide ranks all available tools by empirically measured average score. Structured lookup tools (e.g., `search_mesh_descri ptor` at 18.00, `search_chembl_target` at 17.83) are preferred over text-based search APIs (e.g., `search_uniprot_entity` at 16.44), and both are preferred over direct SPARQL for initial entity resolution.

#### Defensive querying

The guide mandates exploratory queries with `LIMIT_10` to verify structure before attempting comprehensive retrieval, and `LIMI_T`/`OFFSET` chunking for large result sets. Structured predicates from MIE files are required in preference to text search (`bif:contains` or `FILTER(CONTAINS())`) wherever possible. Known hard query patterns (e.g., cross-endpoint joins, specialist databases with sparse data) are explicitly flagged with recommended workarounds.

#### Output quality

The guide requires concise, non-repetitive final answers. Each fact must appear exactly once; meta-commentary (e.g., “Based on my analysis… “) and internal reasoning artifacts are prohibited. Partial answers with explicit acknowledgment of gaps are preferred over incorrect exhaustive attempts.

### Benchmark Question Set and Evaluation

To evaluate TogoMCP, a benchmark of 50 biologically meaningful questions was created following a structured protocol. Questions span five types, with ten per type, and were required to involve at least two RDF Portal databases in 60% of cases and at least three in 20%, with all 23 databases supported at benchmark-creation time covered at least once; the 5 databases added during revision (see section “System Architecture and Environment”) are exposed by TogoMCP but were not retroactively incorporated into the benchmark. Because these 5 databases were not represented during experiment design, the current benchmark metrics should be read as a lower bound on TogoMCP’s effective coverage. As additional databases continue to be integrated into TogoMCP, we plan to extend the benchmark with new questions targeting them and to revise existing questions as needed; this expansion is supported by an ongoing effort to systematize the benchmark question-creation process into a semi-automatic pipeline. Questions recoverable from model pre-training or the published literature were excluded; each candidate was validated against PubMed to confirm that RDF database access was necessary to answer it.

All question bodies are formulated exclusively in natural language; no pre-resolved database identifiers appear in the question text. Answering each question therefore requires the agent to perform entity resolution as part of its workflow: it must call appropriate search APIs to obtain database-specific identifiers before constructing SPARQL queries. This mirrors the two-stage workflow illustrated in Figure 1, where resolving the entity name “TP53” to its ChEMBL target identifier (CHEMBL4096) via `search_chembl_target`—which returns a ranked candidate list, not a single unambiguous hit—is an obligatory intermediate step before any SPARQL can be issued. The same pattern applies across databases: resolving a gene symbol to an NCBI Gene IRI, a disease name to a MONDO or MeSH identifier, or a chemical name to a ChEBI or PubChem compound. Because the agent must select the correct hit from the ranked list, genuine disambiguation is required: the search result for a human gene symbol may include non-human orthologs, splice variants, or—for targets like TP53—both a gene entry and a drug-target entry in different databases. The MIE file for each database includes an `anti_patterns` section that explicitly flags common misresolutions of this kind (e.g., confusing `search_chembl_target` with `search_chembl_molecule`, or selecting a non-human protein when a human isoform is required), ensuring the agent has schema-level guidance for disambiguation at the point of query construction.

The five question types are:

- **Yes/no.** Tests whether a specific entity possesses a binary property (e.g., “Does JAK2 have pathogenic ClinVar variants?”); requires an existence or absence check against the RDF graph using a named entity and a verifiable criterion.
- **Factoid.** Requests a single retrievable value—either a count obtained by comprehensive aggregation (e.g., “How many reviewed human proteins are annotated with kinase activity?”) or a specific attribute lookup (e.g., “What is the EC number of enzyme X?”).
- **List.** Asks for an enumeration of entities satisfying a set of constraints, either comprehensively (“Which proteins have property Y?”) or as a ranked subset (“Top 5 proteins by criterion Z”); the target set size is kept within a verifiable range of 5–100 items.
- **Summary.** Requires multi-dimensional aggregation across two to four facets, drawing on three or more databases, answered in a single synthesized paragraph (e.g., “Characterize the taxonomic distribution, functional categories, and structural coverage of proteins in pathway X”).
- **Choice.** Presents either an explicit bounded list of alternatives (e.g., “Which of [A, B, C, D] has the most X?”) or an unbounded categorical comparison (e.g., “Which taxonomic order has the most proteins with property Y?”), requiring the agent to enumerate and count across all candidate categories before selecting the answer.

Question creation followed a mandatory type-first workflow: the target question type was selected before databases or keywords to enforce balanced type distribution. Biological concepts were grounded in structured vocabularies (GO (6), MONDO, MeSH (5), ChEBI, NCBI Taxonomy) via the OLS4 API (18) before any SPARQL was written, and all descendant terms were retrieved to ensure complete vocabulary coverage. Aggregation queries were verified arithmetically—the sum of per-category counts was checked against an independent `COUN_TDISTINCT` query, with discrepancies treated as coverage gaps requiring correction.

Each question carries two reference-answer fields that together constitute the gold standard. The `exact_answer` field is type-specific: for yes/no, factoid, and choice questions it records the verifiable result extracted directly from executed SPARQL queries at question-creation time, against live database state; for list questions it holds an array of up to ten entries (with database identifiers) drawn from a comprehensively queried and arithmetically verified result set, capped at a verifiable scope of 5–100 items; for summary questions it is intentionally left empty and graded holistically against the accompanying `ideal_answer`. The `ideal_answer` is a single synthesized paragraph written at expert level, integrating quantitative SPARQL results with biological context and containing no meta-references to queries or databases. Benchmark questions were designed to route each factual claim to a single authoritative source, thereby minimising cross-database ambiguity: entity-level facts (e.g., variant pathogenicity classifications) are obtained from their primary curation database (e.g., ClinVar), and cross-database joins rely exclusively on formally declared shared identifiers (e.g., UniProt cross-references to PDB or ChEMBL). When a single entity belongs to multiple categories—causing the per-category sum to exceed the unique entity count—this overlap is documented explicitly in `ideal_answer` (e.g., “8 of 15 proteins participate in more than one inflammasome type”). Question designs in which two databases could return contradictory facts for the same entity are avoided; where such disagreement is structurally possible, the question is rewritten to reference the authoritative source unambiguously. All 50 questions passed an independent QA review that verified coverage completeness, arithmetic consistency of aggregation queries, exhaustive vocabulary coverage, and the absence of circular reasoning.

#### Answers were collected from two agents: a baseline (Claude

Sonnet 4.5 `claude-sonnet-4-5-20250929`, no tools) and a TogoMCP agent (same model with MCP database access, web search explicitly denied). Each question ran in an isolated session with no prior conversation history. Answer quality was then scored by an LLM judge (25) (Claude Opus 4.7) on four criteria (26)—information recall, precision, non-redundancy, and readability—each on a 1–5 scale (total 4–20), enabling direct pairwise comparison between the two agents. Five independent evaluation runs were averaged per question–answer pair to mitigate single-run stochastic variance introduced by the judge’s non-zero generation temperature; the per-question spread of these five runs is reported below in section “Evaluator Reliability and Manual Validation”. To complement the LLM judge with human assessment, a stratified random subset of 20 questions (four per question type) was additionally re-evaluated manually by three domain-expert co-authors. For each selected question, the corresponding baseline and TogoMCP answers were presented in a blinded A/B worksheet with randomized ordering and scored independently on the same four 1–5 dimensions used by the LLM judge; the resulting human scores were compared against the judge’s five-run means on the same subset (section “Evaluator Reliability and Manual Validation” below).

## Results

### Illustrative Examples

Before presenting the systematic evaluation, we briefly illustrate the system’s capabilities with three representative scenarios that motivate the benchmark design.

When provided with the relevant MIE file, the LLM generated SPARQL queries that correctly utilize database-specific biological vocabularies. For instance, given the query “Determine the distribution of human enzymes by their EC number classes using only the reviewed UniProt entries,” the LLM correctly employed the taxonomy IRI `taxonomy/9606` and the Boolean literal `true` for the `up:reviewed` property. Similarly, the MIE files enabled successful use of both standard RDF hierarchical properties (`rdfs:subClassOf*` for GO) and MeSH-specific vocabularies (`meshv:parentTreeNumber`), confirming robust handling of specialized biological ontologies.

The two-stage hybrid approach—API-based entity resolution followed by SPARQL graph exploration—proved essential for cross-database queries. In a complex query seeking human proteins related to cancer pathways with known 3D structures and FDA-approved drug targets, API tools identified 75 unique UniProt entries, compared with only 3 found by a pure SPARQL JOIN approach. This highlighted the failure of SPARQL string filtering for initial entity discovery and motivated the benchmark’s analysis of structured versus text-based search tools.

Finally, the system was tested on a complex hypothesis-generation task: building a model for Alzheimer’s disease progression, integrating multiple RDF Portal datasets (GO, Reactome, UniProt, MONDO, ChEMBL, MeSH). Initially, the LLM hallucinated a non-existent ChEMBL property (`cco:mechanismOfAction`). After provisioning the revised ChEMBL MIE file, the LLM successfully generated correct SPARQL to extract verifiable evidence linking known AD drugs (e.g., donepezil) to their target proteins. This example illustrates the mechanism behind the ablation study’s central finding: MIE schema files inject precise, database-specific structural knowledge that the LLM cannot reliably reproduce from its pre-training alone, and removing them measurably reduces system performance (quantified below).

These three scenarios—schema-faithful vocabulary use, hybrid-workflow leverage on cross-database queries, and elimination of hallucinated predicates—instantiate the three mechanisms that the 50-question benchmark systematically evaluates next.

### Benchmark Evaluation

To systematically quantify the system’s utility, 50 biologically grounded questions spanning five types (yes/no, factoid, list, summary, choice; 10 each) were evaluated by comparing the TogoMCP agent against a no-tool baseline. Each answer was independently scored five times by an LLM judge (25) (Claude Opus 4.7; a fixed model snapshot was used for all conditions to avoid evaluator drift) across four dimensions (26)—recall, precision, non-redundancy, and readability—on a 1–5 scale (total 4–20), yielding 250 question–run pairs per condition. In addition to the full system (**With Guide**), three ablation conditions were tested: exclusion of the Usage Guide, but with an explicit instruction to call `find_databases` and `get_MIE_file` before queries (**MIE-Instr**); exclusion of the Usage Guide with no such instruction (**No-Instr**); and exclusion of the `get_MIE_file` tool entirely (**No MIE**).

#### Overall Performance

TogoMCP delivered a statistically significant and practically large improvement over the baseline. Across 250 question–run pairs the mean total score rose from 15.10 to 18.55 (Δ = +3.45, Cohen’s *d* = 1.82, Wilcoxon *p* < 0.001; Table 2). TogoMCP won on 94.4% of individual evaluations, tied on 4.0%, and lost on only 1.6%. The baseline never achieved a perfect score of 20; TogoMCP reached it on 107 of 250 evaluations (42.8%), spanning 26 distinct questions. The gains were strongly concentrated in *information recall* (+2.23 on the 5-point scale), confirming that tool-augmented access to live RDF databases supplies factual content that the LLM alone cannot fabricate. Precision also improved substantially (+1.04), and both non-redundancy and readability showed marginal positive gains (+0.11 and +0.07, respectively).

**Table 2.** Overall score comparison (mean ± SD, 50 questions × 5 evaluation runs).

| | Baseline | TogoMCP | $\Delta$ |
| --- | --- | --- | --- |
| Total | 15.10 ± 1.51 | 18.55 ± 1.65 | +3.45 |
| Recall | 1.85 | 4.08 | +2.23 |
| Precision | 3.61 | 4.65 | +1.04 |
| Non-redundancy | 4.80 | 4.91 | +0.11 |
| Readability | 4.84 | 4.91 | +0.07 |

#### Performance by Question Type

The benefit of TogoMCP varied markedly across question types (Table 3). Yes/no questions showed the largest gain (Δ = +4.22, 100% win rate), followed by factoid (+3.82, 96%) and list (+3.58, 90%)—types with precise, verifiable answers where database evidence is decisive. Choice questions also benefited substantially (+3.20, 96%). Summary questions saw the smallest gain (+2.44, 90% win rate, 4% loss rate), reflecting the greater challenge of multi-dimensional synthesis where additional factual content does not always offset the verbosity introduced by multi-step tool-use responses.

**Table 3.** Score improvement (Δ) and win/loss rates by question type.

| Type | Baseline | TogoMCP | $\Delta$ | Win / Lose |
| --- | --- | --- | --- | --- |
| yes/no | 15.62 | 19.84 | +4.22 | 100% / 0% |
| factoid | 13.82 | 17.64 | +3.82 | 96% / 0% |
| list | 14.30 | 17.88 | +3.58 | 90% / 4% |
| choice | 16.22 | 19.42 | +3.20 | 96% / 0% |
| summary | 15.52 | 17.96 | +2.44 | 90% / 4% |

Across all 250 evaluations, TogoMCP lost on only 1.6% of pairs. Only one question showed a net negative average delta: Q021 (summary), which asks for a landscape characterization of human proteasome subunits across GO cellular component subcomplexes—a multi-hop aggregation task where the baseline’s concise hedged response outscored the more verbose tool-augmented answer. One list question (Q044) broke even on average. These marginal failures represent cases where query complexity exceeds the benefit of structured database access within the evaluation’s readability and non-redundancy dimensions.

#### Tool Usage and Workflow Efficiency

A negative correlation was observed between the number of tool calls and the TogoMCP score (Spearman *ρ* = *−*0.44, Pearson *r* = *−*0.28). The optimal range was 5–9 tool calls (mean score 19.1), with 15–19 calls dropping to 16.7. A small number of high-volume workflows (30+ calls) still reached near-perfect scores, indicating that excessive chaining is most often the model’s response to early failure rather than an intrinsic ceiling. A similar pattern held for `run_sparql` invocations: questions with 0–2 SPARQL calls averaged 19.2–20.0, while those with ten or more calls fell to 17.1.

Tool effectiveness varied across categories. Structured lookup tools returning precise identifiers—`search_mesh_descr iptor` (mean score 20.00, *n* = 4 questions), `ncbi_esearch` (19.25, *n* = 13), and `search_reactome_entity` (18.60, *n* = 3)— outperformed text-based entity-search APIs (`search_uniprot_entity` at 17.79, *n* = 16; `search_chembl_target` at 17.70, *n* = 4). The gap of approximately 1–2 points is modest in absolute terms, but the directional preference for structured lookup is consistent across the categories.

Workflow-pattern analysis showed that 72% of evaluations followed the full Guide-prescribed sequence (`find_database s` *→* `get_MIE_file` *→* search *→* `run_sparql`; mean score 18.5). A further 22% completed entity discovery via MIE content alone and proceeded directly to SPARQL without an explicit search step (mean 18.6), and a smaller 6% subset answered the question via NCBI E-utilities alone, with no SPARQL call to the RDF Portal (mean 19.2); these latter cases involve questions whose required content can be retrieved through structured NCBI APIs without invoking the SPARQL endpoint. Procedural compliance with the Usage Guide was high: 100% initial compliance with the `TogoMCP_Usage_Guide` call, 94% MIE-file reading, and 78% search-tool usage.

The mean latency was 137 s (17.4× baseline) and the mean cost was $0.38 per question (70× baseline). Although substantial, these overheads are justified for precision-critical queries where the baseline cannot retrieve the requested information: the 15 questions with the largest improvements (Δ ≥ 5) averaged a 12.7× time ratio, whereas the 10 worst-performing questions averaged 21.7×.

#### Ablation Study: MIE Files and the Usage Guide

To dissect the contributions of the system’s two key components—the MIE schema files and the Usage Guide protocol—four experimental conditions were compared (Table 4).

**Table 4.** Four-condition ablation comparison. All metrics are computed against each condition’s own no-tool baseline. MIE-Instr: no Usage Guide, but explicit instruction to call find databases and get MIE file. No-Instr: no Usage Guide, no instructions. No MIE: get MIE file excluded entirely.

|  | With Guide | MIE-Instr | No-Instr | No MIE |
| --- | --- | --- | --- | --- |
| $\Delta$ vs. baseline | +3.45 | +3.46 | +3.22 | +2.96 |
| Cohen’s $d$ | 1.82 | 1.56 | 1.19 | 1.38 |
| $p$-value | < 0.001 | < 0.001 | < 0.001 | < 0.001 |
| Win rate | 94.4% | 92.8% | 81.2% | 83.2% |
| Perfect (=20) | 42.8% | 48.4% | 40.0% | 33.6% |
| Tools/question | 12.4 | 12.1 | 20.9 | 16.9 |
| Cost/question | \$0.39 | \$0.38 | \$0.41 | \$0.39 |

##### All conditions outperform baseline; MIE files are the primary contributor to the mean

All four ablation conditions delivered statistically significant and large improvements over the no-tool baseline (*d* = 1.19–1.82, all *p* < 0.001), demonstrating that the system as a whole is robust to component removal. To isolate individual component contributions, we compared per-question improvement scores (Δ = TogoMCP *−* baseline) across conditions using paired Wilcoxon signed-rank tests. Removing `get_MIE_file` reduced the improvement from Δ = +3.45 to Δ = +2.96—a 0.50-point reduction that is marginally non-significant by a two-sided test (Wilcoxon *p* = 0.067) but whose 90% bootstrap CI [+0.04, +0.94] excludes zero. Cohen’s *d* on the per-question paired difference is 0.25, a small effect at the question level. The No MIE condition achieved this performance by substituting iterative schema exploration for the MIE file: `find_databases` (called on 100% of questions, per the Usage Guide) identified the relevant endpoint, `get_graph_list` supplied the named-graph URI needed for the mandatory `FROM` clause, and iterative `run_sparql` calls (median 5 per question, with the most complex cases reaching ~15– 19) empirically discovered predicates and class structures. This strategy succeeds for most databases but falls short on databases with non-standard RDF structures (e.g., AMR Portal blank-node phenotypes, BacDive organism hierarchies) where the MIE file’s annotated examples and anti-patterns provide irreplaceable guidance.

##### The MIE instruction captures the full benefit of the Usage Guide *on average*

The MIE-Instr condition—identical to No-Instr except for a single system-prompt instruction to call `find_databases` and `get_MIE_file` before queries—recovered virtually the entire benefit of the full Usage Guide on the mean per-question improvement (Δ = +3.46 vs. +3.45; two-sided Wilcoxon *p* = 0.877; 95% bootstrap CI [*−*0.40, +0.41]). By contrast, the No-Instr condition, lacking both the Usage Guide and explicit instruction, spontaneously called `get_MIE_file` on only 18% of questions and `find_databases` on only 10%, yet still achieved Δ = +3.22 (two-sided Wilcoxon *p* = 0.739 vs. With Guide). The extra tool calls in No-Instr (20.9 vs. 12.4 per question) reflect compensatory trial-and-error SPARQL rather than structured schema discovery. A plausible reading is that the MIE instruction acts as an implicit early-exit cue: once the agent has loaded concrete worked examples from the MIE file, the empirically observed need for prolonged schema-exploration SPARQL is reduced, and the additional procedural rules in the Usage Guide (workflow ordering, explicit SPARQL budgets, interleaving warnings) confer no measurable mean-score gain beyond what the MIE instruction alone already produces.

##### Component value hierarchy and the Usage Guide’s worst-case role

The pairwise comparisons above (With Guide vs. MIE-Instr indistinguishable; No-Instr vs. No MIE also indistinguishable, *p* = 0.269) locate the only significant separation between MIE-equipped and No MIE conditions. Mean comparisons do not, however, fully capture the Usage Guide’s contribution. The Guide also narrows the spread of outcomes and reduces downside risk: across 250 individual evaluations, With Guide produced a loss (TogoMCP score below baseline) on only 4 evaluations (1.6%), significantly fewer than MIE-Instr (12 losses, 4.8%; Fisher exact one-sided *p* = 0.036), No-Instr (15 losses, 6.0%; *p* = 0.009), or No MIE (10 losses, 4.0%; *p* = 0.087); see Supplementary Table S2. The between-question SD of mean Δ across 50 questions is 1.90 for With Guide versus 2.21 for MIE-Instr, 2.71 for No-Instr (Levene *p* = 0.005 vs. With Guide), and 2.14 for No MIE. Two complementary worst-case statistics tell a consistent story. First, the minimum per-question mean Δ across the 50 questions is *−*0.20 for With Guide, versus *−*1.20, *−*3.20, and *−*2.40 for MIE-Instr, No-Instr, and No MIE respectively. Second, when worst-case is defined at the evaluator-run level—taking the minimum TogoMCP score across the five evaluator runs for each question, then comparing the resulting per-question worst-case scores in a paired Wilcoxon test—With Guide stochastically dominates No-Instr (mean difference +0.78, *p* = 0.013) but is statistically indistinguishable from MIE-Instr (mean difference +0.18, *p* = 0.21). Taken together, the availability of `get_MIE_file` is the primary source of marginal gain in the mean, and a single instruction to use it is the highest-leverage intervention for mean-score performance. The Usage Guide adds a further— but smaller—reduction in downside risk relative to MIE-Instr, most clearly visible as a roughly threefold reduction in the per-evaluation loss rate. When neither the Usage Guide nor an MIE instruction is provided, both the mean and the variance degrade in tandem.

### Evaluator Reliability and Manual Validation

#### Stochastic variation across five evaluation runs

The LLM judge (Opus 4.7) operates at non-zero generation temperature, so independent evaluations of the same question– answer pair are not deterministic; averaging five independent runs per pair is the manuscript’s design choice for absorbing this evaluator-side noise. The per-question SD across the five runs quantifies how large that noise is. For the With Guide condition the mean per-question evaluator SD was 0.46 (median 0.50, max 1.30) on TogoMCP totals and 0.94 (median 0.89, max 1.79) on baseline totals, on the 4–20 score scale; 17 of 50 TogoMCP questions had identical totals across all five runs. Across all four ablation conditions pooled (n = 400 condition×question×arm cells), the mean evaluator SD was 0.79. Averaging across the five evaluator runs per cell reduces this raw evaluator noise to approximately 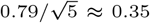 well below both the between-question SD of the mean Δ across 50 questions (1.90) and the across-condition Δ gap (~0.5 points). The standard error of the mean Δ across the 50 questions is therefore approximately 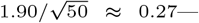the appropriate scale against which to interpret the +3.45-point With Guide improvement. Evaluator stability varies systematically by question type: yes/no and choice questions have the lowest mean per-question Δ-SD across the five runs (0.74 and 0.66 respectively), summary questions an intermediate value (0.99), and factoid and list questions the highest between-run variance (1.41 and 1.22), reflecting the larger information surface on which the judge can disagree.

#### Manual validation by three domain-expert co-authors

To provide a human reference against which to interpret the LLM judge, a stratified random subset of 20 questions (four per question type) was re-evaluated manually by three domain-expert co-authors. Each annotator completed all 20 blinded A/B pairs independently, without inspecting the LLM-judge output. Each annotator’s mean Δ confirms the system’s benefit with high statistical significance, but the absolute magnitude of the improvement varies between annotators (Table 5): all three identify a large effect with Cohen’s *d* ranging from 0.90 to 1.52, all *p ≤* 0.002, and a consistent direction of improvement (win/tie/loss combined: 47/4/9 across the 60 question-annotator pairs). The single composite reference obtained by averaging the three annotators’ scores per question yields a manual baseline total of 11.28 ± 3.15, a manual TogoMCP total of 16.40 ± 2.08, and a manual Δ of +5.12 ± 3.82 (paired Cohen’s *d* = 1.34, Wilcoxon *p* = 2.0 × 10*^−4^*; win/tie/loss 18/1/1). On the same 20 questions, the LLM judge’s five-run mean Δ was +3.48 ± 2.15 (*d* = 1.62; win/tie/loss 18/1/1). All three independent human annotators thus identified absolute improvements larger than the LLM judge but in the same effect-size class and with the same statistical conclusion: TogoMCP delivers a significant, large improvement over the no-tool baseline.

**Table 5.** Per-annotator and composite manual evaluation on the 20-question subset (vs. their respective baseline answer on the same question), compared with the LLM judge’s five-run mean on the same subset.

| Reference | Baseline | TogoMCP | $\Delta$ | Cohen’s $d$ |
| --- | --- | --- | --- | --- |
| Annotator A | 10.25 ± 3.51 | 16.80 ± 2.91 | +6.55 ± 4.31 | 1.52 |
| Annotator B | 11.65 ± 5.02 | 17.35 ± 1.98 | +5.70 ± 5.83 | 0.98 |
| Annotator C | 11.95 ± 1.99 | 15.05 ± 2.86 | +3.10 ± 3.46 | 0.90 |
| Mean of three | 11.28 ± 3.15 | 16.40 ± 2.08 | +5.12 ± 3.82 | 1.34 |
| LLM judge (5-run mean) | 14.89 ± 1.59 | 18.37 ± 2.07 | +3.48 ± 2.15 | 1.62 |

#### Inter-annotator agreement

Reliability across the three independent human annotators was moderate at the level of individual ratings and substantially higher for the composite (mean-of-three) reference, the configuration used in the subsequent LLM-judge comparison. On the 40 individual score pairs (20 questions × 2 answers each), the single-rater intraclass correlation (36) was ICC(2,1) = 0.675 (“moderate” by the Koo–Li benchmarks (37)), rising to ICC(2,*k*=3) = 0.862 (“good”) for the average of the three raters; Krippendorff’s *α* (interval) on the same 40 pairs was 0.670. On the 20 per-question Δ values the agreement was lower, ICC(2,1) = 0.461 and ICC(2,*k*=3) = 0.719 (*α* = 0.432), reflecting the higher variance that arises from differencing two noisier ratings. Pairwise Pearson correlations on totals were *r* = 0.771 (A–B), 0.707 (A–C), and 0.675 (B–C); pairwise sign-agreement on per-question Δ was 13– 15 out of 20. The inter-annotator gap is attributable mainly to a baseline-leniency difference: Annotator B rated baseline answers +1.40 points higher than Annotator A on average, while their TogoMCP scores differed by only +0.55; Annotator C used a narrower overall score range ([9, 19] versus [5, 20] for Annotators A and B) and identified four questions on which TogoMCP underperformed (versus zero for Annotator A and five for Annotator B). Despite these individual-rater differences, the average-of-three reference is statistically reliable (ICC(2,*k*=3) = 0.862 on totals, 0.719 on per-question Δ), justifying its use as a composite human baseline against which to compare the LLM judge.

#### Agreement between manual and LLM-judge scores

Across the 40 individual score pairs, the Pearson correlation between the mean-of-three manual scores and the LLM-judge totals was *r* = 0.908 (*p* < 10*^−^*15; Spearman *ρ* = 0.902); restricted to the 20 per-question Δ values, *r* = 0.765 (*p* = 8.3 × 10*^−5^*; Spearman *ρ* = 0.817). Individual annotator correlations with the LLM judge bracket this composite result: *r* on totals = 0.873 (A), 0.747 (B), and 0.869 (C); *r* on per-question Δ = 0.768 (A), 0.476 (B), and 0.775 (C). The direction of the per-question Δ (TogoMCP wins, ties, or loses) agreed between the mean-of-three reference and the LLM judge on 16 of 20 questions; sign-agreement on the same comparison for individual annotators was 17/20 (A), 13/20 (B), and 15/20 (C). Per-dimension agreement was strongest on Recall (mean-of-three vs. LLM *r* = 0.750 on per-question Δ; mean Δ +1.80 vs. +2.39 on the 1–5 scale) and Precision (*r* = 0.729; +1.63 vs. +1.00). The two style dimensions showed substantially lower agreement: Non-redundancy (*r* = 0.426; mean Δ +1.12 vs. +0.07) and Readability (*r* = 0.044; mean Δ +0.57 vs. +0.02). The LLM judge thus tracks the human consensus closely on Recall and Precision—the two dimensions that carry the bulk of the measured improvement—while systematically attenuating the human-perceived advantage of TogoMCP on the style dimensions. The three-annotator pass therefore supports the LLM-judge ranking on the dimensions where the system’s benefit is concentrated, and bounds the direction of bias on the dimensions where it is not.

## Discussion

The benchmark results show that an LLM augmented with MCP-mediated tools can substantially outperform an unaided LLM for natural language querying of RDF knowledge graphs (Δ = +3.45, Cohen’s *d* = 1.82), and that this improvement is robust across all tested component configurations. We organize the discussion around four themes: the role of schema context, the dynamics of the two-stage hybrid workflow, practical cost– performance trade-offs, and current limitations with directions for future work.

### Schema context as the primary marginal contributor

The ablation study reveals that all four experimental conditions—With Guide, MIE-Instr, No-Instr, and No MIE— deliver large, statistically significant improvements over the no-tool baseline (*d* = 1.19–1.82, all *p* < 0.001), demonstrating that TogoMCP’s benefits are robust to component removal. Within this picture, MIE files emerge as the primary source of marginal gain: the No MIE condition partially compensates through iterative schema exploration—using `get_graph_list` to obtain the named-graph URI required by mandatory `FROM` clauses, then empirically discovering predicates through repeated `run_sparql` calls (per-question counts and the resulting deficit reported in section “Ablation Study: MIE Files and the Usage Guide”)—but this strategy is insufficient for databases whose RDF structures cannot be inferred from endpoint introspection alone (e.g., blank-node phenotypes in AMR Portal, organism hierarchies in BacDive).

Providing MIE files together with a single system-prompt instruction to use them (the MIE-Instr condition) recovered the full benefit of the elaborate Usage Guide protocol on the mean per-question improvement (Δ = +3.46 vs. +3.45; two-sided Wilcoxon *p* = 0.877 for the difference; 95% bootstrap CI [*−*0.40, +0.41]). The Usage Guide’s additional guidance on workflow ordering, SPARQL budgets, and interleaving warnings contributed no measurable improvement on the mean beyond what the MIE instruction alone conferred. It does, however, narrow the spread of outcomes: with the Usage Guide the per-evaluation loss rate falls to 1.6% (4/250), versus 4.8% under MIE-Instr (Fisher exact one-sided *p* = 0.036) and 6.0% under No-Instr (*p* = 0.009), and the between-question SD of mean Δ contracts from 2.71 (No-Instr) and 2.21 (MIE-Instr) to 1.90 (With Guide; Levene *p* = 0.005 vs. No-Instr). A plausible reading is that the MIE instruction acts as an implicit early-exit cue—once the agent has loaded concrete worked examples from the MIE file, the need for prolonged exploratory SPARQL is reduced—while the Usage Guide’s procedural rules add a secondary insurance against the rarer pathological workflows that survive the MIE-instruction route. This result has a practical implication: for similar tool-augmented LLM systems (12; 27; 28), investing in high-quality, concise schema documentation delivered at query time is likely 1.62 to be far more impactful than investing in complex behavioral protocols.

The success of MIE files can be understood as a form of dynamic in-context learning. Rather than relying on the LLM’s pre-trained (and often inaccurate) knowledge of RDF vocabularies, MIE files inject precise, database-specific structural information—shape expressions, example queries, anti-patterns—directly into the context window at the point of need, as illustrated by the `cco:mechanismOfAction` example in section “Results”. This finding is consistent with broader observations in the LLM literature that retrieval-augmented approaches outperform pure parametric memory for tasks requiring domain-specific factual knowledge (35).

A qualitative inspection of per-question score differences across conditions suggests that two MIE sections carry the irreplaceable residual value: `sparql_query_examples` (concrete working queries the LLM can adapt; 21% of MIE content) and `cross_database_queries` (multi-graph query idioms for co-querying databases that share the same SPARQL endpoint, e.g. UniProt + Rhea on the SIB endpoint; 9%). Both lack any architectural substitute outside the MIE file itself. By contrast, `shape_expressions` (correct predicate names and IRI patterns) is partially substituted by `get_graph_list` (whose system-graph filtering and database-name relevance ranking are described in section “System Architecture and Environment”) plus trial-and-error SPARQL iteration, and `schema_info` is consumed server-side by `find_databases` before the LLM ever reads it.

This subcomponent ranking is corroborated by two patterns at the question-type and database levels. By question type, multi-database synthesis tasks—summary (Δ_WG_ *−* Δ_NoMIE_ = +0.70) and choice (+1.48)—show the largest residual No MIE deficit, whereas factoid (*−*0.52) and list (*−*0.40) types, where direct retrieval substitutes are sufficient, show No MIE matching or exceeding With Guide. By database, the largest With Guide advantages over No MIE occur on questions targeting BacDive (Δ = +5.4), PubTator (Δ = +3.6–+5.0), and complex AMR Portal phenotype queries (Δ = +2.8), all of which require niche database idioms or schema details not reachable through endpoint introspection within the SPARQL budget. The two With Guide failure cases (questions 044 and 021), discussed in detail under section “Performance variation across question types”, reflect entity-resolution and response-style errors rather than schema-vocabulary gaps that MIE context could address. A formal subcomponent ablation is left as a dedicated future study.

These findings resonate with, and extend, recent related work. SPARQL-LLM (13), the most closely related system, also relies on ShEx schemas and example queries but retrieves them dynamically via embedding similarity from a vector index, whereas TogoMCP delivers a curated, per-database MIE document in its entirety. Our ablation result—that removing MIE files produces a modest but consistently-directed reduction in per-question improvement (two-sided *p* = 0.067, with 90% bootstrap CI excluding zero)—suggests that comprehensive, editorially reviewed schema context may be more effective than similarity-retrieved fragments, at the cost of manual curation effort. Rangel et al. (11) independently observed that semantic clues in SPARQL (meaningful variable names, inline comments) boost fine-tuned model performance by 33%; the anti-pattern sections and annotated query examples in MIE files serve an analogous role in the zero-shot setting, steering the LLM away from hallucinated predicates without any training. Ali et al. (15) confirmed the same principle in a clinical setting: an ontology-grounded knowledge graph eliminated hallucinations that standalone LLMs produced, paralleling the contrast between our With MIE and No MIE conditions. Taken together, these converging results across fine-tuning, RAG, and tool-augmented paradigms reinforce the general principle that explicit structural context—whether injected via training data enrichment, retrieved schema fragments, or curated MIE files—is the primary determinant of LLM accuracy on knowledge-graph queries.

### Workflow ordering and the two-stage hybrid approach

The intended two-stage workflow—entity resolution via external APIs followed by SPARQL graph exploration—is the protocol the Usage Guide prescribes, and the evaluation runs show that the system follows it on the majority of questions (workflow-pattern statistics reported in section “Tool Usage and Workflow Efficiency”). The short-circuit paths (MIE-only entity discovery, or NCBI-E-utilities-only retrieval) score equally well or slightly better than the full pipeline—an artefact of question difficulty distribution, since the easier questions are typically the ones for which the MIE file or a structured NCBI lookup is self-sufficient—but the practical implication holds: as MIE files are enriched with more comprehensive entity IRIs and example patterns, the search stage can become less critical, reducing both latency and the risk of compounding errors.

Within the search stage itself, structured lookup tools outperformed text-based entity-search APIs by approximately 1–2 points on the 4–20 scale (per-tool means reported in section “Tool Usage and Workflow Efficiency”). The gap is modest but the directional preference for structured ID resolution over free-text search is consistent across all tested categories, and future protocol iterations should continue to prioritize structured lookup wherever it is available.

### Tool call economy and cost–performance trade-offs

A consistent finding across all conditions was the negative correlation between tool call volume and answer quality (Spearman *ρ* = *−*0.44, Pearson *r* = *−*0.28). The optimal range was 5–9 tool calls per question (mean score 19.1); beyond 15 calls, scores fell to 16.7. A similar pattern held for SPARQL invocations: questions with 0–2 `run_sparql` calls averaged 19.2–20.0, whereas those issuing ten or more dropped to 17.1. Excessive tool chaining does not reflect thoroughness but rather the system struggling to recover from an incorrect initial approach—a behavior most pronounced in the **No-Instr** condition, where 48% of questions required 18 or more tool calls (versus 14% under **With Guide**).

The practical costs are non-trivial: mean latency of 137 s (17.4× baseline) and the mean cost of $0.38 per question (70× baseline). However, these averages obscure an important pattern. The 15 questions with the largest improvements (Δ ≥ 5) averaged a 12.7× time ratio, while the 10 worst-performing questions averaged 21.7×. The cost–benefit ratio is most favorable precisely for the queries where TogoMCP provides the greatest value—those requiring database-specific factual content that the LLM alone cannot produce. The current Usage Guide directly addresses this with an implemented early-exit rule: upon reaching two consecutive SPARQL calls without convergence, the protocol mandates halting and either simplifying the query, switching to a search tool, or synthesizing from partial results—rather than continuing to accumulate cost.

### Performance variation across question types

The benefit of TogoMCP was strongly modulated by question type. Yes/no questions showed the largest gain (Δ = +4.22, 100% win rate), followed by factoid (+3.82, 96%/0%), list (+3.58, 90%/4%), and choice (+3.20, 96%/0%) questions— types with precise, verifiable answers, consistent with the system’s strength in retrieving specific facts from structured databases. Summary questions showed the smallest gain (+2.44, 90% win rate, 4% loss rate), reflecting the difficulty of multi-dimensional synthesis across several databases, where each integration step introduces opportunities for error accumulation and where the baseline’s hedged, general-knowledge answers can remain competitive despite lacking direct database evidence. The reduction relative to other types is now modest, however: even on summary questions TogoMCP wins on 90% of evaluations.

The set of questions on which TogoMCP underperformed has shrunk to two: question 044 (a list question on glycan classification, Δ = 0.0; vocabulary mismatch between the GlyCosmos and GlyConnect curation scopes) and question 021 (a summary question on the proteasome landscape, Δ = *−*0.2; evaluator-penalized verbosity). Both failure modes are addressable: question 044 requires richer cross-vocabulary mapping (for example, inclusion of GlyConnect predicates in the GlyCosmos MIE entries), and question 021 reflects an interaction between the model’s multi-database synthesis behaviour and the LLM judge’s preference for concise prose, rather than a deficiency in the SPARQL retrieval itself.

The five benchmark question types—yes/no, factoid, list, summary, and choice—represent the core of factual biological querying but do not exhaust the range of queries that life-science researchers direct at RDF knowledge graphs in practice. Three further query patterns deserve particular attention. *Hypothesis-generation queries* ask the system to integrate multi-database evidence to support or refute a biological model: for example, mapping disease-associated proteins to their curated pathways and approved drug targets to assess mechanistic coherence. *Multi-hop cross-database reasoning* chains identifier conversions across three or more endpoints in a single session—a pattern qualitatively more demanding than the two-hop cross-endpoint workflows that already account for the ten worst benchmark failures. *Negative-existence queries* ask the system to confirm that a specific combination of properties does *not* appear in any curated database entry—requiring exhaustive enumeration rather than selective retrieval.

A detailed illustration of the first two patterns is provided in Supplementary File 1: a multi-scale Alzheimer’s disease analysis session in which TogoMCP anchors the disease to standardized identifiers across MONDO, MeSH, and DOID; retrieves Swiss-Prot-reviewed AD-associated proteins from UniProt via SPARQL; converts accessions to PDB, NCBI Gene, and ChEMBL target identifiers via `togoid_convertId`; queries Reactome for curated CDK5 and amyloid-aggregation pathways; and retrieves approved therapeutics from ChEMBL—spanning six databases across four SPARQL endpoints in a single coherent session. These open-ended, multi-step session types were excluded from the formal benchmark because their scope makes LLM-judge scoring less reliable than for the bounded question types evaluated here; the 50-question benchmark therefore provides a conservative lower bound on the system’s performance across the full breadth of life-science queries researchers are likely to pose.

### Limitations

Several limitations should be acknowledged. First, the principal evaluation channel is an LLM judge (Claude Opus 4.7) rather than human expert assessment, which may introduce systematic biases. A three-annotator manual validation on a 20-question subset (see section “Evaluator Reliability and Manual Validation”) shows strong rank-order agreement between the mean-of-three human reference and the LLM judge (Pearson *r* = 0.908 on individual totals, *r* = 0.765 on per-question Δ, with direction agreement on 16 of 20 questions); the LLM judge tracks the human consensus closely on Recall and Precision—the two dimensions that carry the bulk of the measured improvement—while substantially attenuating the human-perceived advantage of TogoMCP on the style dimensions of non-redundancy and readability.

Three caveats temper this conclusion. (i) Reliability across the three annotators was moderate at the single-rater level (ICC(2,1) = 0.675 on totals, 0.461 on per-question Δ) and rose to “good” only after averaging the three raters (ICC(2,*k*=3) = 0.862 and 0.719 respectively), reflecting genuine individual differences in scoring calibration that are absorbed but not eliminated by the composite reference. (ii) Three of the five question types (yes/no, choice, list) require the annotator to integrate three long text artefacts (the ideal answer and two candidate answers) per question, a cognitive load that risks rewarding superficially fluent responses; this pattern is visible in the smaller relative Δ on yes/no and choice questions for the most lenient annotator. (iii) Annotators reported greater difficulty in calibrating Recall and Precision on choice and list questions, where the expected answer surface varies more between questions than on yes/no or factoid types, and greater confidence on questions within their own primary research domains than on those outside it. Each of these factors limits the resolution of the manual validation, but none reverses its direction: three independent expert annotators each identified a large, statistically significant improvement, and the LLM judge ranks questions in the same order as the human consensus.

Second, the benchmark of 50 questions, while carefully designed to span five question types and the 23 databases supported at the time of benchmark creation, represents a limited sample of the possible query space; characterising the 5 databases added during revision through the benchmark remains future work.

Third, the system was evaluated exclusively with the Claude family (Sonnet 4.5 and Opus 4.7); generalizability to other model families remains untested. The MIE file format is plain-text YAML with no Claude-specific features, and the core mechanism—delivering structured schema context at inference time—is consistent with gains reported in fine-tuning (11) and RAG-based (13) paradigms, suggesting the approach is not inherently model-family-dependent; systematic cross-model evaluation is a committed direction for future work.

Fourth, the MIE files were created semi-automatically with manual review, and their quality directly determines system performance—scaling this process to additional databases will require more automated validation pipelines.

Fifth, the benchmark deliberately avoids entities for which name-to-identifier resolution would be ambiguous even for domain experts—cases where two curators might legitimately disagree on the correct database entry, such as polysemous gene symbols with conflicting cross-species annotations or chemical names with multiple structurally distinct database entries. Benchmark questions were instead designed to have a single defensible correct resolution, so that evaluation scores reflect SPARQL reasoning quality rather than unresolvable naming ambiguity. Edge cases requiring more nuanced handling of such conflicts are reserved for future evaluation.

Finally, the current system does not exploit cross-endpoint JOIN queries natively. The 28 supported databases are distributed across eight SPARQL endpoints; although these endpoints do technically support SPARQL 1.1 `SERVICE` federation, our preliminary testing found such federated queries to be highly unreliable in practice—frequent timeouts, intermittent failures, and unpredictable performance even on superficially simple joins—making them unsuitable as a production-quality query mechanism. This operational hurdle is a concrete instance of a broader systemic challenge in distributed bioinformatics: the FAIR principles (38) call for data to be Interoperable across resources, but in practice Interoperability at the level of a single, federated query is difficult to guarantee even when each constituent endpoint is independently FAIR-compliant, since federation reliability depends on infrastructure and protocol behaviour that lie outside any individual database’s control. TogoMCP’s sequential bridging strategy is therefore best understood as a pragmatic workaround for a standards-compliance gap rather than a permanent substitute for reliable federation. TogoMCP therefore answers questions that span databases on different endpoints through a sequential multi-query workflow: the agent retrieves identifiers from one endpoint and passes them as a `VALUES` clause to a subsequent query on a second endpoint. This introduces a specific error-propagation risk: if the first query returns an incomplete identifier set—for example, because a cross-reference link (`rdfs:seeAlso`) is absent from a UniProt entry, or because TogoID has no mapping for a particular entity—the downstream query receives a silently truncated `VALUES` clause and returns a plausible but incorrect count, with no error raised to alert the user.

As a concrete example, one benchmark question asks which human chromosome harbours the most cardiomyopathy-associated protein-coding genes. Answering it requires the agent to retrieve all reviewed UniProt proteins annotated with cardiomyopathy from the SIB endpoint, extract their NCBI Gene cross-references via `rdfs:seeAlso` links, and pass those gene identifiers to the NCBI endpoint for chromosomal counting. Any UniProt entry lacking a curated `rdfs:seeAlso` link to NCBI Gene is silently excluded from the downstream count, potentially misidentifying the leading chromosome.

Quantitatively, 42 of the 50 benchmark questions (84%) involve databases on different SPARQL endpoints and therefore depend on this sequential bridging pattern. `togoid_convertId` was invoked in 9 of 50 questions (18% of the benchmark); its mean score (18.27) is slightly below the overall mean (18.55), and the two worst-performing questions (q044 and q021, discussed in section “Performance variation across question types”) both involve cross-database integration challenges.

Improving the reliability of the sequential bridging pattern through pre-flight identifier coverage checks and intermediate result validation is a priority for future system hardening.

### Future directions

Several avenues for improvement possibly follow from the results. Enriching MIE files with more comprehensive entity IRI inventories and common query templates could reduce dependence on the search stage, improving both latency and accuracy. Implementing automatic workflow restart upon detection of the reversed (SPARQL *→* search) pattern could eliminate a significant source of poor performance. The current Usage Guide already encodes early-exit and question-type routing rules; future work could expose these as explicit software-level controls to guarantee adherence independent of the LLM’s behavioral compliance. Evaluation with human domain experts would strengthen confidence in the system’s biological utility. Long-term sustainability of the MIE file collection is itself a data-governance concern: as underlying ontologies and endpoint schemas evolve, a MIE file can silently drift out of sync with the live RDF structure it describes. At present, updates to the RDF Portal are detected through periodic manual review, and the corresponding MIE files are revised accordingly; automating this update-detection step is a priority for future work, for example via a monitoring pipeline that periodically reissues each MIE file’s `sparql_query_examples` against the live endpoint and flags query failures or schema-shape mismatches for expert review, extending the validation already performed at onboarding (section “Context Provisioning via MIE Files”) into a continuous, scheduled process rather than a periodic manual one. Finally, as MCP adoption grows, interoperability with additional MCP servers (e.g., for clinical trial data) could extend the system’s coverage without architectural changes.

## Conclusion

This study demonstrates that a natural-language cross-database query system for the RDF Portal is feasible when the LLM is treated as a protocol-driven inference engine orchestrating specialized tools via MCP. The system achieved a large improvement over an unaided baseline (Cohen’s *d* = 1.82), with gains concentrated in information recall and precision, and win rates exceeding 80% for question types with precise, verifiable answers. The ablation study found that all component configurations deliver significant improvements, with MIE schema files as the primary marginal contributor to the mean and a one-line instruction to use them recovering the same mean-score benefit as the elaborate behavioural protocol; the procedural protocol contributes a smaller but complementary benefit by narrowing outcome variance and reducing the per-evaluation loss rate roughly threefold.

These findings point to a general design principle for tool-augmented LLM systems over structured knowledge bases: concise, dynamically delivered schema context is the highest-leverage component for mean performance, while procedural orchestration plays a distinct, complementary role in managing the worst case. Combined with the two-stage hybrid workflow —structured entity resolution followed by schema-guided SPARQL—this architecture substantially lowers the barrier to querying complex biological knowledge graphs, providing researchers with a transparent and verifiable platform for evidence-based discovery. Future work will focus on enriching the MIE file coverage, validating the system with human domain experts, and extending interoperability to additional MCP-connected data sources.

## Supplementary data

**Supplementary File 1** is a multi-scale Alzheimer’s disease analysis document generated using TogoMCP, illustrating hypothesis-generation and multi-hop cross-database reasoning across six databases and four SPARQL endpoints.

**Supplementary Table S1** provides per-question score deltas across all four ablation conditions for all 50 benchmark questions (section “Ablation Study: MIE Files and the Usage Guide”, Table 4).

**Supplementary Table S2** provides the distribution of per-question Δ values across the four conditions, the pairwise statistical tests (Wilcoxon signed-rank, Fisher exact loss-rate, Levene variance) with bootstrap confidence intervals, and the per-condition loss summary with the identities of all net-loss questions (section “Ablation Study: MIE Files and the Usage Guide”).

## Author contributions statement

A.R.K. and Y.Y. conceived the experiments. A.R.K. and

Y.Y. implemented the TogoMCP system. A.R.K. and S.B.L. implemented the benchmark evaluation scripts. A.R.K. conducted the experiments. A.R.K., Y.Y., and T.F. performed the manual evaluation of the 20-question subset. A.R.K., Y.Y., S.B.L., T.F., and J.E.L.G. analysed the results. A.R.K. and

Y.Y. wrote the manuscript. A.R.K., Y.Y., S.B.L., T.F., and

J.E.L.G. reviewed the manuscript.

## Acknowledgment

A.R.K. thanks Anastasios Nentidis for helping create BioASQ-inspired (26) QA sets, and Shuichi Kawashima, Toshiaki Katayama, Issaku Yamada, Jin-Dong Kim, Yuki Moriya, and Yuna Oikawa for valuable feedback. We also acknowledge and thank the participants of the DBCLS BioHackathon 2025 (BH25), with whom several of the authors had fruitful discussions and gained deeper insights into this work.

## Data availability

The remote TogoMCP server is available at https://togomcp.rdfportal.org/. The source code, including the benchmark questions and results, used in this work is available at Zenodo: https://doi.org/10.5281/zenodo.20076385 (the latest code at https://github.com/dbcls/togomcp).

## Footnotes

1 https://modelcontextprotocol.io/specification/2025-11-25

2 Namespace prefixes used in this paper: `up:` = http://purl.uniprot.org/core/ (UniProt Core Ontology); cco: = http://rdf.ebi.ac.uk/terms/chembl/ (ChEMBL Core Ontology); meshv: = http://id.nlm.nih.gov/mesh/vocab# (MeSH RDF Vocabulary); rdfs: = http://www.w3.org/2000/01/rdf-schema# (RDF Schema, W3C standard); bif: = Virtuoso built-in function prefix (OpenLink Virtuoso SPARQL engine).

