## Supplementary File 1 for "TogoMCP: Natural Language Querying of Life-Science Knowledge Graphs via Schema-Guided LLMs and the Model Context Protocol"

### ALZHEIMER'S DISEASE

#### Multi-Scale Pathophysiology Analysis

*Database-Integrated Analysis: SPARQL · TogoID · UniProt · Reactome · ChEMBL · PubMed*
Generated: March 2026

#### Executive Summary

Alzheimer's disease (AD) is the most prevalent neurodegenerative disorder worldwide, characterized by progressive cognitive decline and dementia. It is defined pathologically by extracellular deposition of amyloid-β (Aβ) peptides into senile plaques and intracellular accumulation of hyperphosphorylated tau (pTau) as neurofibrillary tangles (NFTs). The disease operates across multiple biological scales: amyloidogenic processing of the amyloid precursor protein (APP) by BACE1 and γ-secretase (PSEN1/2) drives Aβ42 overproduction at the molecular level; CDK5 deregulation and tau hyperphosphorylation disrupt cytoskeletal integrity at the cellular level; neuroinflammation mediated by microglia and astrocytes accelerates damage at the tissue level; and progressive hippocampal and cortical atrophy produces the hallmark clinical syndrome of memory loss and cognitive impairment. The APOE ε4 allele remains the strongest genetic risk factor for sporadic AD. Approved therapies include acetylcholinesterase inhibitors (donepezil, galantamine, rivastigmine), NMDA receptor antagonism (memantine), and the newly approved anti-amyloid monoclonal antibody lecanemab.

#### 1. Disease Ontology Cross-References

The following table shows cross-database identifier mappings retrieved via TogoID and OLS4:

| Source DB | Source ID | Target DB | Target ID |
|---|---|---|---|
| MONDO | MONDO:0004975 | MeSH | D000544 |
| MONDO | MONDO:0004975 | DOID | DOID:10652 |
| MONDO | MONDO:0004975 | HP (phenotype) | HP:0002511 |
| SNOMED | 26929004 | NCIt | NCIT:C2866 |
| EFO | EFO:0000249 | DOID (via EFO) | DOID:10652 |

DOID:10652 definition (OLS4): 'A tauopathy characterized by memory lapses, confusion, emotional instability and progressive loss of mental ability… marked histologically by neurofibrillary tangles and plaques containing beta-amyloid.'

### 2. Molecular Level

#### 2.1 Core AD Proteins (UniProt SPARQL Query Results)

SPARQL queries against UniProt RDF (Swiss-Prot reviewed, human: taxonomy/9606) identified the following core proteins. Disease annotations were retrieved using bif:contains full-text search.

| UniProt ID | Mnemonic | Protein Name | Gene | Disease Role / SPARQL Annotation |
|---|---|---|---|---|
| P05067 | A4_HUMAN | Amyloid precursor protein | APP | Processed by BACE1 + γ-secretase → Aβ peptides; mutations cause familial AD |
| P49768 | PSN1_HUMAN | Presenilin-1 | PSEN1 | Catalytic subunit of γ-secretase; >200 FAD mutations increase Aβ42:Aβ40 ratio |
| P49810 | PSN2_HUMAN | Presenilin-2 | PSEN2 | Second γ-secretase catalytic subunit; FAD mutations modulate Aβ42 production |
| P56817 | BACE1_HUMAN | Beta-secretase 1 | BACE1 | β-site APP cleaving enzyme; rate-limiting step in amyloidogenic processing; major drug target |
| P10636 | TAU_HUMAN | Microtubule-associated protein tau | MAPT | SPARQL: 'neuronal cytoskeleton progressively disrupted → paired helical filaments (PHF)'; reduced O-GlcNAcylation → hyperphosphorylation |
| P02649 | APOE_HUMAN | Apolipoprotein E | APOE | SPARQL: 'APOE*4 allele risk increased from 20% to 90%… mean age at onset decreased 84→68 years'; impairs Aβ clearance; disrupts blood-brain barrier |

#### 2.2 Additional Disease-Associated Proteins (SPARQL – UniProt)

Full-text SPARQL bif:contains query on 'Alzheimer' across disease annotations returned additional proteins:

| UniProt ID | Mnemonic | Protein Name | AD Relevance (SPARQL) |
|---|---|---|---|
| P10636 | TAU_HUMAN | Microtubule-associated protein tau | PHF-TAU / AD P-TAU in NFTs; corticobasal degeneration overlap |
| P37840 | SYUA_HUMAN | Alpha-synuclein | Major non-Aβ component of AD amyloid plaque; forms Lewy body inclusions |
| O00429 | DNM1L_HUMAN | Dynamin-1-like protein (DRP1) | Aβ-induced S-nitrosylation triggers excessive mitochondrial fission, synaptic loss |

| | | | |
|---|---|---|---|
| O15294 | OGT1_HUMAN | O-GlcNAc transferase (OGT) | Reduced O-GlcNAcylation of MAPT/TAU → increased phosphorylation in AD brain |
| Q9NRA0 | SPHK2_HUMAN | Sphingosine kinase 2 | Nuclear localization in AD brains; cytosolic expression correlates inversely with amyloid plaque density |
| P21554 | CNR1_HUMAN | Cannabinoid receptor 1 | CNR1 heterozygous deletion → decreased PSD95, accelerated memory impairment in AD mouse models |

#### 2.3 Cross-Database ID Mapping (TogoID)

TogoID conversions linking core AD proteins across six databases (route: uniprot → ncbigene, pdb, chembl_target):

| UniProt | Gene | NCBI Gene | Protein | PDB Structures (selected) | ChEMBL Target | PDB Count |
|---|---|---|---|---|---|---|
| P05067 | APP | 351 | APP | 1AAP, 2BEG, 3BAE, 6GFI, 7Q4B… | CHEMBL2487 | >200 |
| P49768 | PSEN1 | 5663 | PSEN1 | 2KR6, 5A63, 6IDF, 6LQG, 8X52… | CHEMBL2473 | 25+ |
| P49810 | PSEN2 | 5664 | PSEN2 | 7Y5T, 7Y5X, 7Y5Z | CHEMBL3708 | 3 |
| P56817 | BACE1 | 23621 | BACE1 | 1FKN, 1M4H, 2B8L, 4FRI, 5I3V… | CHEMBL4822 | >400 |
| P10636 | MAPT | 4137 | TAU | 1I8H, 6TJO, 7QJV, 8Q27, 9GG0… | CHEMBL1293224 | >200 |
| P02649 | APOE | 348 | APOE | 1B68, 1LPE, 2KC3, 7FCR, 8GRX… | N/A | 30+ |

### 3. Pathway Level

#### 3.1 Dysregulated Pathways (Reactome SPARQL Query Results)

SPARQL queries against the Reactome BioPAX RDF graph (FROM <http://rdf.ebi.ac.uk/dataset/reactome>) identified the following key AD pathways:

| Reactome Pathway (SPARQL Result) | Mechanistic Summary |
|---|---|
| Deregulated CDK5 triggers multiple neurodegenerative pathways in AD models | Aβ → $Ca^{2+}$ dysregulation → calpain cleaves p35 → p25; CDK5:p25 hyperactivates CDC25A/B/C → CDK1/2/4 activation → neuronal death |
| Amyloid fiber formation (R-HSA-624) | Aggregation pathway for misfolded proteins including Aβ42 and TAU; AGRN:beta-amyloid fibril interaction via GAG chains confirmed by SPARQL |

| | |
|---|---|
| APP processing reactions (Reactome BioPAX) | SPARQL confirmed: HTRA2 degrades APP; AP4 binds APP; AGRN binds Beta amyloid fibril (UniProt O00468 AGRIN) |
| Cholinergic neurotransmission (inferred) | Basal forebrain cholinergic neuron loss; target of AChE inhibitor drug class |
| APOE-dependent lipid transport in CNS (UniProt SPARQL) | APOE*4: impairs Aβ clearance, disrupts BBB, activates MAP3K12/AP-1 → upregulates APP transcription (UniProt annotation) |

##### 3.2 CDK5 Pathway Detail (SPARQL Comment Text)

The Reactome SPARQL query returned a rich mechanistic description: neurotoxic insults (Aβ, ischemia, oxidative stress) disrupt intracellular $Ca^{2+}$ homeostasis → calpain cleaves p35 → p25. The CDK5:p25 complex translocates to cytoplasm and nucleus, phosphorylating CDC25A (at S40, S116, S261), CDC25B (S50, T69, S160, S321, S470) and CDC25C (T48, T67, S122, T130, S168, S214), releasing them from 14-3-3 inhibition, which in turn activates CDK1, CDK2, and CDK4 kinases leading to neuronal death. Higher CDC25A/B/C activities were confirmed in human AD clinical samples vs. age-matched controls.

### 4. Cellular Level

Multiple cell types are directly implicated:

| Cell Type | Dysfunction Mechanism | Evidence Source |
|---|---|---|
| Pyramidal neurons | NFT accumulation → cytoskeletal disruption; CDK5:p25 → aberrant cell cycle entry → apoptosis | MAPT/TAU SPARQL (UniProt P10636); Reactome CDK5 pathway |
| Basal forebrain cholinergic neurons | Selective early degeneration; loss of ACh synthesis capacity → cognitive impairment | Drug mechanism basis for AChE inhibitors |
| Microglia | Aβ triggers microglial activation → inflammatory cytokine release → NLRP3 inflammasome; TREM2/PLCG2 variants alter microglial Aβ clearance | PLCG2 (P16885) found in Reactome SPARQL APP-processing reactions |
| Astrocytes | Reactive astrogliosis around plaques; impaired glutamate uptake; GFAP elevation (AD biomarker) | Literature/PubMed review (Liu et al. 2024) |
| Oligodendrocytes | Transition to disease-associated state; dual role in myelin defense and disease progression; active participation in amyloid/tau pathology | Kedia & Simons 2025, Nature Neuroscience, DOI:10.1038/s41593-025-01873-x |
| Mitochondria (all neurons) | DRP1 S-nitrosylation (DNM1L, UniProt O00429) → excessive fission → synaptic loss; impaired OXPHOS | UniProt SPARQL disease annotation (O00429) |

### 5. Tissue and Organ Level

AD affects the brain in a stereotyped spatiotemporal pattern. The hippocampus and entorhinal cortex are affected earliest, manifesting as episodic memory failure; the disease then spreads to association cortices, parietal lobes, and eventually primary sensorimotor areas. Tau pathology spreads following Braak staging (I–VI). Amyloid PET and CSF/plasma Aβ42:Aβ40 ratios now enable preclinical detection decades before symptoms.

| Region | Histopathology | Clinical Correlate |
|---|---|---|
| Entorhinal cortex / Hippocampus | Earliest NFT deposition (Braak I–II); neuronal loss in CA1, subiculum | Episodic memory failure; inability to form new memories |
| Parietal / temporal association cortex | Diffuse and neuritic Aβ plaques; synaptic pruning; NFT spread (Braak III–IV) | Language deficits, apraxia, visuospatial impairment |
| Prefrontal cortex | Later-stage NFT and neuronal loss; reduced glucose metabolism on FDG-PET | Executive dysfunction, personality change |
| Basal forebrain / locus coeruleus | Cholinergic neuron loss; noradrenergic degeneration | Attention, arousal, and memory consolidation failure |
| Cerebrovasculature / BBB | APOE4 disrupts BBB (pericyte loss); cerebral amyloid angiopathy (CAA); VEGF dysregulation | Cerebrovascular disease contribution to cognitive decline |

### 6. Clinical Level

#### 6.1 Disease Staging & Biomarkers

| Stage | Clinical Features | Biomarkers |
|---|---|---|
| Preclinical (A+T-N-) | Asymptomatic; detectable Aβ deposition 15–20 years before symptoms | Aβ-PET+, CSF Aβ42↓, Aβ42:Aβ40 ratio↓ |
| MCI due to AD (A+T+N-) | Subtle memory decline; preserved ADL function | p-Tau181/217↑, CSF t-Tau↑, hippocampal atrophy on MRI |
| Mild AD (A+T+N+) | Episodic memory loss, language difficulties, disorientation | Tau-PET+, GFAP↑, NfL↑ in plasma/CSF |
| Moderate AD | Executive dysfunction, behavioral changes, impaired ADLs | Extensive cortical tau spread; FDG-PET hypometabolism |
| Severe AD | Profound cognitive/physical decline; loss of speech, continence, ambulation | Widespread cortical atrophy; global synaptic loss |

#### 6.2 HP Phenotype Mapping (TogoID: MONDO → HP)

TogoID conversion (MONDO:0004975 → hp_phenotype) returned HP:0002511 (Alzheimer disease). Key associated HP phenotypes include:

| HP Term | Clinical Phenotype Description |
|---|---|
| HP:0002511 | Alzheimer disease (primary phenotype) |
| HP:0000726 | Dementia — progressive decline in multiple cognitive domains |
| HP:0002354 | Memory impairment — specifically episodic memory failure |
| HP:0001300 | Parkinsonism — occurs in some late-stage AD cases |
| HP:0002145 | Frontotemporal dementia — overlap in some MAPT-related cases |

### 7. Treatment Mechanisms

#### 7.1 Drug-Target Relationships (ChEMBL + TogoID)

| Drug Name | ChEMBL ID | PubChem CID | DrugBank | Target | Mechanism of Action |
|---|---|---|---|---|---|
| Donepezil | CHEMBL502 | 3152 | DB00843 | AChE/ BChE | Reversible AChE inhibitor → ↑synaptic ACh → compensates for cholinergic neuron loss |
| Galantamine | CHEMBL659 | 9651 | DB00674 | AChE / nAChR | AChE inhibitor + allosteric nAChR modulator (sensitization) |
| Rivastigmine | CHEMBL636 | 77991 | DB00989 | AChE/ BChE | Pseudo-irreversible dual AChE/BChE inhibitor; available as transdermal patch |
| Memantine | CHEMBL807 | 4054 | DB01043 | NMDA-R | Low-affinity uncompetitive NMDA receptor antagonist → prevents glutamate excitotoxicity |
| Lecanemab | CHEMBL3833321 | N/A | N/A | Aβ protofibrils | Humanized IgG1 mAb targeting Aβ protofibrils; FDA-approved 2023 (disease-modifying) |
| Aducanumab | — | — | — | Aβ fibrils/plaques | Anti-Aβ mAb (accelerated FDA approval 2021); targets aggregated Aβ |

#### 7.2 Drug Target Proteins (UniProt/ChEMBL Mapping)

| Drug | ChEMBL Target | UniProt | Target Protein | Rationale |
|---|---|---|---|---|
| Donepezil/ | CHEMBL220 | P22303 (ACHE) | Acetylcholinesterase | Compensates basal |

| | | | | |
|---|---|---|---|---|
| Galantamine/ Rivastigmine | | | | forebrain cholinergic neuron loss |
| Memantine | CHEMBL1907631 | Q05586 (GRIN1) | NMDA receptor GluN1 | Reduces excitotoxic $Ca^{2+}$ influx from chronic low-level glutamate |
| BACE1 inhibitors (e.g., verubecestat) | CHEMBL4822 | P56817 (BACE1) | Beta-secretase 1 | Block rate-limiting step in Aβ generation; >400 crystal structures in PDB |
| Lecanemab | — | P05067 (APP) | Aβ protofibrils (from APP) | Clears soluble Aβ protofibrils; slows cognitive decline in trials |

### 8. Integrated Multi-Scale Disease Model

The causal cascade from molecular defects to clinical symptoms follows this hierarchy, grounded in database evidence:

| Scale | Key Events (Database-Derived) | Causal Link to Next Scale |
|---|---|---|
| MOLECULAR | APP (P05067) processed by BACE1 (P56817) + PSEN1/2 γ-secretase → Aβ42 overproduction; MAPT (P10636) hyperphosphorylated by CDK5/GSK3β; APOE4 (P02649) impairs Aβ clearance | Aβ42 aggregates into protofibrils → full plaques; p-tau forms NFTs |
| PATHWAY | Reactome: CDK5:p25 phosphorylates CDC25A/B/C → CDK1/2/4 activation; Amyloid fiber formation pathway (R-HSA-624); APOE4 activates MAP3K12/AP-1 → upregulates APP transcription (UniProt SPARQL) | Aberrant cell cycle re-entry → mitotic catastrophe; synaptic Aβ oligomers disrupt LTP |
| CELLULAR | Pyramidal neuron cytoskeletal collapse (TAU PHF); mitochondrial fragmentation via DRP1 S-nitrosylation (O00429); cholinergic neuron loss; microglial Aβ phagocytosis failure; OGT (O15294) → reduced O-GlcNAcylation → ↑tau phosphorylation | Synaptic loss precedes neuronal death; neuroinflammation amplifies damage |
| TISSUE / ORGAN | Hippocampal and entorhinal cortex atrophy (earliest); Braak stage I–VI tau propagation; CAA in cerebrovasculature; BBB disruption (APOE4-pericyte loss); glucose hypometabolism on FDG-PET | Volume loss on MRI; white matter disruption (oligodendrocyte dysfunction) |
| CLINICAL | DOID:10652, MeSH:D000544, HP:0002511; progressive memory loss → language → executive function → global impairment; 5–10 year survival after diagnosis | Clinical staging (CDR, MMSE); biomarker-defined A/T/N framework |

### 9. Key Literature Evidence (PubMed)

Based on articles retrieved from PubMed:

| Citation | Key Findings Relevant to This Analysis |
|---|---|
| Hampel H et al. Mol Psychiatry 2021;26:5481-5503. DOI: 10.1038/s41380-021-01249-0 | Systematic review establishing the Aβ pathway as the core biochemical hallmark of AD; supports distinct interactions of Aβ species with tau, neuroinflammation, and neurochemical imbalance; basis for Aβ-targeting therapies |
| Serrano-Pozo A et al. Lancet Neurol 2021;20:68-80. DOI: 10.1016/S1474-4422(20)30412-9 | APOE4 risk increase from 20%→90% with allele dose confirmed; APOE pathogenesis now extends to tau degeneration, microglial/astrocyte responses, and BBB disruption beyond Aβ-centric mechanisms |
| Liu E, Zhang Y, Wang JZ. Transl Neurodegener 2024;13:45. DOI: 10.1186/s40035-024-00432-x | Comprehensive 2024 update: risk factors (APOE variants, infections), molecular mechanisms, newly approved Aβ vaccines (lecanemab), novel tau-targeting strategies, peripheral biomarkers |
| Kedia S, Simons M. Nat Neurosci 2025;28:446-456. DOI: 10.1038/s41593-025-01873-x | Emerging 2025 evidence: oligodendrocytes transition to disease-associated state in AD; dual role in immune modulation, myelin defense and disease progression; important for non-neuronal disease biology |

### 10. Data Sources and Tools

| Tool / Database | Version / Endpoint | Data Retrieved |
|---|---|---|
| UniProt RDF (SPARQL) | https://rdfportal.org/sib/sparql | Disease-annotated proteins, GO terms, function annotations (bif:contains, Swiss-Prot reviewed, taxonomy/9606) |
| Reactome RDF (SPARQL) | https://rdfportal.org/ebi/sparql | AD pathways (bp:Pathway), biochemical reactions, protein participants with UniProt xrefs |
| TogoID API | https://togoid.dbcls.jp/ | ID conversions: uniprot→ncbigene, pdb, chembl_target; chembl_compound→pubchem, drugbank; mondo→mesh, doid, hp |
| OLS4 (EBI) | https://www.ebi.ac.uk/ols4/ | DOID:10652 definition; MONDO, SNOMED, NCIT, EFO cross-references |
| ChEMBL (search API) | https://www.ebi.ac.uk/chembl/ | Drug molecule search: donepezil, galantamine, rivastigmine, memantine, lecanemab |
| PubMed (MCP) | https://pubmed.ncbi.nlm.nih.gov/ | Literature review: 4 key references (PMID 33340485, 34456336, 39232848, 39881195) |

Note: All SPARQL queries used bif:contains for full-text search with up:reviewed 1 filter on UniProt. Reactome queries targeted the named graph <http://rdf.ebi.ac.uk/dataset/reactome>. TogoID conversions used the togoid_convertId API with explicit source,target routes. Literature citations retrieved via PubMed MCP server and properly attributed per PubMed terms of use.

### Appendix: Session Log

The following is an annotated log of the TogoMCP session that produced this document, describing the databases queried, tools invoked, and key findings at each phase.

#### Overview

This session was a structured bioinformatics analysis using TogoMCP to integrate multiple life-science databases and produce a multi-scale pathophysiology report on Alzheimer's disease (AD).

*This session was generated by following the disease analysis workflow prompt at https://github.com/dbcls/togomcp/blob/main/workflows/disease_analysis.md with the input parameter "disease name: Alzheimer's disease".*

#### Phase 1 — Disease Ontology Mapping

Using TogoID and OLS4, the primary AD concept was mapped across ontologies:

• MONDO:0004975 → MeSH D000544, DOID 10652, HP HP:0002511

• Cross-references: SNOMED 26929004, NCIT C2866, EFO 0000249

#### Phase 2 — Protein Discovery via SPARQL (UniProt)

SPARQL queries against the UniProt RDF endpoint retrieved Swiss-Prot reviewed human proteins annotated with Alzheimer's disease:

| UniProt ID | Symbol | Role |
|---|---|---|
| P05067 | APP | Amyloid precursor protein |
| P49768 | PSEN1 | γ-secretase catalytic subunit |
| P56817 | BACE1 | Rate-limiting β-secretase, major drug target |
| P10636 | MAPT | Tau — forms neurofibrillary tangles when hyperphosphorylated |
| P02649 | APOE | APOE4 allele impairs Aβ clearance, disrupts BBB |

Additional proteins linked to mitochondrial fission (DRP1), tau O-GlcNAcylation (OGT1), and cannabinoid signaling (CNR1) were also retrieved.

#### Phase 3 — Cross-Database ID Conversion (TogoID)

UniProt accessions were converted to identifiers in downstream databases:

• UniProt → NCBI Gene: e.g., P05067 → Gene ID 351

• UniProt → PDB: revealed hundreds of structural entries per protein

• UniProt → ChEMBL Target: e.g., BACE1 → CHEMBL4822

#### Phase 4 — Pathway Analysis via SPARQL (Reactome)

Key AD-associated pathways identified:

• Deregulated CDK5 pathway: Aβ → $Ca^{2+}$ dysregulation → calpain cleaves p35 to p25 → CDK5:p25 phosphorylates CDC25A/B/C → aberrant CDK1/2/4 activation → neuronal death

• Amyloid fiber formation (R-HSA-624): aggregation pathway for Aβ42 and tau

#### Phase 5 — Drug-Target Analysis (ChEMBL + TogoID)

Approved AD therapeutics retrieved from ChEMBL:

| Drug | Mechanism | ChEMBL ID |
|---|---|---|
| Donepezil | AChE inhibitor | CHEMBL502 |
| Galantamine | AChE/nAChR modulator | CHEMBL807 |
| Rivastigmine | AChE/BChE inhibitor | CHEMBL659 |
| Memantine | NMDA antagonist | CHEMBL636 |
| Lecanemab | Anti-Aβ protofibril mAb (FDA-approved 2023) | CHEMBL3833321 |

#### Phase 6 — Literature Evidence (PubMed)

Key recent reviews retrieved via PubMed MCP:

• Serrano-Pozo et al., Lancet Neurol 2021 — APOE genetics and pathophysiology
• Hampel et al., Mol Psychiatry 2021 — Amyloid-β pathway systematic review
• Liu et al., Transl Neurodegener 2024 — AD mechanisms to therapies
• Kedia & Simons, Nat Neurosci 2025 — Oligodendrocytes in AD

#### Tools and Databases Used

| Tool / Database | Purpose |
|---|---|
| OLS4 | Disease ontology lookup |
| TogoID | Cross-database identifier conversion |
| UniProt SPARQL | Protein–disease annotations |
| Reactome SPARQL | Biological pathway data |
| ChEMBL | Drug and target data |
| PubMed MCP | Literature retrieval |
