## Supplementary Table S1 for "TogoMCP: Natural Language Querying of Life-Science Knowledge Graphs via Schema-Guided LLMs and the Model Context Protocol"

Per-question score deltas across ablation conditions
TogoMCP: Natural Language Querying of Life-Science Knowledge Graphs via Schema-Guided LLMs and the Model Context Protocol

This table accompanies the main manuscript (section "Ablation Study: Contributions of MIE Files and the Usage Guide", Table 2). All scores are means over five evaluation runs using an LLM judge (Claude Opus 4.7) on a 4–20 scale. Δ values are TogoMCP score minus baseline score for each ablation condition. Bold Δ values indicate gains ≥2.0; red values indicate negative deltas (TogoMCP underperformed baseline). Conditions: WG = With Guide (full system); MIE-Instr (no Usage Guide, explicit MIE instruction); No-Instr (no Usage Guide, no instruction); No MIE (`get_MIE_file` excluded).

Table 1: Per-question score deltas across all four ablation conditions. Scores are averages over five evaluation runs (LLM judge, 4–20 scale). Δ values are TogoMCP minus baseline for each condition. Type abbreviations: Y/N = yes/no, Fact = factoid, List = list, Sum = summary, Cho = choice.

| **Q** | **Type** | **Baseline** | WG | ΔWG | ΔMIE-Instr | ΔNo-Instr | ΔNo MIE |
|---|---|---|---|---|---|---|---|
| Q001 | Y/N | 18.0 | 20.0 | **+2.0** | +1.6 | **+2.2** | +0.8 |
| Q007 | Y/N | 13.4 | 19.8 | **+6.4** | **+7.6** | **+7.8** | **+6.0** |
| Q012 | Y/N | 15.0 | 19.4 | **+4.4** | **+6.4** | **+3.0** | **+3.6** |
| Q017 | Y/N | 16.8 | 19.8 | **+3.0** | **+2.4** | -3.2 | -2.4 |
| Q020 | Y/N | 13.8 | 20.0 | **+6.2** | **+6.6** | **+6.8** | **+6.6** |
| Q026 | Y/N | 17.0 | 20.0 | **+3.0** | **+2.8** | **+2.4** | **+2.2** |
| Q032 | Y/N | 17.2 | 20.0 | **+2.8** | **+3.4** | +0.6 | +0.2 |
| Q036 | Y/N | 14.0 | 20.0 | **+6.0** | **+6.4** | **+6.6** | **+5.6** |
| Q042 | Y/N | 16.4 | 19.4 | **+3.0** | **+3.2** | **+2.0** | **+4.0** |
| Q046 | Y/N | 14.6 | 20.0 | **+5.4** | **+6.0** | **+5.0** | **+3.4** |
| Q002 | Fact | 14.0 | 17.8 | **+3.8** | **+3.2** | **+3.0** | **+4.4** |
| Q003 | Fact | 13.8 | 16.8 | **+3.0** | **+5.4** | **+7.2** | **+2.6** |
| Q006 | Fact | 13.8 | 16.8 | **+3.0** | +1.4 | **+3.4** | **+3.4** |
| Q011 | Fact | 13.8 | 16.8 | **+3.0** | **+2.0** | **+5.2** | **+4.8** |
| Q014 | Fact | 13.8 | 19.6 | **+5.8** | **+5.4** | **+7.0** | **+6.6** |
| Q022 | Fact | 13.8 | 16.8 | **+3.0** | **+2.8** | **+4.6** | **+4.4** |
| Q027 | Fact | 13.8 | 15.8 | **+2.0** | +1.4 | **+2.6** | **+5.0** |
| Q031 | Fact | 13.8 | 16.0 | **+2.2** | **+4.2** | **+2.6** | **+2.8** |
| Q043 | Fact | 13.8 | 20.0 | **+6.2** | **+5.8** | **+6.8** | **+6.6** |
| Q047 | Fact | 13.8 | 20.0 | **+6.2** | **+5.8** | **+3.0** | **+2.8** |
| Q009 | List | 15.0 | 16.2 | +1.2 | **+2.0** | +0.4 | **+2.8** |
| Q013 | List | 16.2 | 18.6 | **+2.4** | -1.2 | +1.8 | **+3.2** |
| Q018 | List | 14.2 | 19.2 | **+5.0** | **+6.2** | **+5.8** | **+5.2** |
| Q023 | List | 14.0 | 17.6 | **+3.6** | **+3.6** | **+6.2** | **+2.6** |
| Q028 | List | 14.0 | 20.0 | **+6.0** | **+6.2** | **+6.2** | **+6.8** |
| Q033 | List | 14.0 | 17.6 | **+3.6** | **+3.2** | **+6.4** | **+5.4** |
| Q039 | List | 14.0 | 20.0 | **+6.0** | **+5.8** | **+6.2** | **+3.0** |
| Q040 | List | 15.0 | 18.8 | **+3.8** | **+2.0** | **+5.2** | **+3.6** |
| Q044 | List | 12.8 | 12.8 | +0.0 | **+2.6** | +0.6 | **+5.0** |
| Q048 | List | 13.8 | 18.0 | **+4.2** | **+3.6** | **+4.0** | **+2.2** |
| Q004 | Sum | 16.0 | 17.0 | +1.0 | +1.8 | +1.4 | +0.0 |
| Q008 | Sum | 15.2 | 17.2 | **+2.0** | **+4.4** | +0.0 | +0.4 |
| Q015 | Sum | 15.2 | 16.8 | +1.6 | **+4.0** | -0.8 | +1.2 |
| Q021 | Sum | 15.0 | 14.8 | -0.2 | **+2.6** | **+2.0** | **+3.0** |

*continued on next page*

Table 1 – *continued*

| **Q** | **Type** | **Baseline** | WG | ΔWG | ΔMIE-Instr | ΔNo-Instr | ΔNo MIE |
|---|---|---|---|---|---|---|---|
| Q024 | Sum | 16.4 | 19.6 | **+3.2** | +1.2 | **+3.8** | **+3.2** |
| Q029 | Sum | 14.8 | 20.0 | **+5.2** | **+4.4** | **+3.4** | **+3.2** |
| Q034 | Sum | 14.8 | 18.0 | **+3.2** | **+4.4** | +0.0 | -0.2 |
| Q037 | Sum | 14.8 | 18.2 | **+3.4** | **+3.2** | -1.0 | **+2.2** |
| Q045 | Sum | 17.2 | 18.4 | +1.2 | +1.2 | +0.4 | +1.2 |
| Q049 | Sum | 15.8 | 19.6 | **+3.8** | **+3.6** | **+3.4** | **+3.2** |
| Q005 | Cho | 18.0 | 20.0 | **+2.0** | +0.0 | +1.0 | +0.2 |
| Q010 | Cho | 14.2 | 20.0 | **+5.8** | **+7.2** | **+6.8** | **+5.0** |
| Q016 | Cho | 17.6 | 18.6 | +1.0 | +0.4 | **+6.6** | +0.6 |
| Q019 | Cho | 13.8 | 20.0 | **+6.2** | **+6.0** | +0.2 | +1.0 |
| Q025 | Cho | 18.0 | 19.0 | +1.0 | +1.6 | +1.0 | +1.2 |
| Q030 | Cho | 17.0 | 18.2 | +1.2 | -1.0 | +0.0 | +1.0 |
| Q035 | Cho | 17.8 | 19.0 | +1.2 | +0.8 | +1.6 | +0.2 |
| Q038 | Cho | 14.0 | 20.0 | **+6.0** | +1.6 | +1.4 | +1.2 |
| Q041 | Cho | 17.8 | 19.4 | +1.6 | +1.2 | +1.4 | +1.2 |
| Q050 | Cho | 14.0 | 20.0 | **+6.0** | **+6.4** | **+7.0** | **+5.6** |
| **Mean** | | **15.10** | **18.55** | **+3.45** | **+3.46** | **+3.22** | **+2.96** |
