## Supplementary Table S2 for "TogoMCP: Natural Language Querying of Life-Science Knowledge Graphs via Schema-Guided LLMs and the Model Context Protocol"

### Aggregate distribution and pairwise statistical tests of ablation conditions
### TogoMCP: Natural Language Querying of Life-Science Knowledge Graphs via Schema-Guided LLMs and the Model Context Protocol

This table accompanies the main manuscript (section “Ablation Study: Contributions of MIE Files and the Usage Guide”, sub-paragraphs “All conditions outperform baseline; MIE files are the primary contributor to the mean”, “The MIE instruction captures the full benefit of the Usage Guide *on average*”, and “Component value hierarchy and the Usage Guide’s worst-case role”). All scores are means over five evaluation runs using an LLM judge (Claude Opus 4.7) on a 4–20 scale. Per-question Δ is the TogoMCP score minus baseline score, computed within each condition’s matched no-tool run.

**Conditions:** WG = With Guide (full system); MIE-Instr = no Usage Guide, but explicit instruction to call `find_databases` and `get_MIE_file`; No-Instr = no Usage Guide, no instructions; No MIE = `get_MIE_file` excluded entirely.

**Panel A. Distribution of per-question mean Δ (n = 50 per condition).**

| Condition | min | $Q_{10}$ | $Q_{25}$ | median | mean | $Q_{75}$ | $Q_{90}$ | max | SD |
|---|---|---|---|---|---|---|---|---|---|
| WG | −0.20 | +1.18 | +2.00 | +3.10 | +3.45 | +5.35 | +6.02 | +6.40 | 1.90 |
| MIE-Instr | −1.20 | +1.16 | +1.65 | +3.20 | +3.46 | +5.70 | +6.40 | +7.60 | 2.21 |
| No-Instr | −3.20 | +0.00 | +1.10 | +3.00 | +3.22 | +6.10 | +6.80 | +7.80 | 2.71 |
| No MIE | −2.40 | +0.20 | +1.20 | +3.00 | +2.96 | +4.70 | +5.64 | +6.80 | 2.14 |

Means match the corresponding row of Table 2 in the main manuscript (within rounding). Between-question SD is computed across the 50 per-question mean Δ values for each condition.

**Panel B. Pairwise statistical tests, With Guide as reference.**

| Comparison vs. WG (other condition) | Mean Δ gap (WG − other) | Wilcoxon $p$ (two-sided) | Fisher loss $p$ (one-sided) | Levene SD $p$ (WG vs. other) |
|---|---|---|---|---|
| MIE-Instr | −0.004 | 0.877 | 0.036 | 0.265 |
| No-Instr | +0.232 | 0.739 | 0.009 | 0.005 |
| No MIE | +0.496 | 0.067 | 0.087 | 0.565 |

*Mean Δ gap*: difference in mean per-question Δ between With Guide and the comparison condition; positive values indicate that With Guide produced a larger improvement on average. *Wilcoxon p*: paired Wilcoxon signed-rank test on the 50 per-question mean Δ values (two-sided). *Fisher loss p*: Fisher’s exact one-sided test on the 2×2 contingency of loss (TogoMCP score below baseline) vs. no-loss evaluations across the 250 question–run pairs per condition; tests whether With Guide produces fewer losses than the comparison condition. *Levene SD p*: Levene test for equality of between-question SD (50 per-question mean Δ values) under center=mean. Worst-case Wilcoxon $p$ values (paired comparison of per-question worst-case TogoMCP score across five evaluator runs) are reported inline in the main manuscript and are not repeated here.

**Panel B (continued). Bootstrap confidence intervals for the paired difference (WG − other), n = 20,000 resamples of the 50 per-question paired differences.**

| Comparison vs. WG | 95% CI | 90% CI |
|---|---|---|
| MIE-Instr | [−0.40, +0.41] | [−0.34, +0.34] |
| No-Instr | [−0.38, +0.85] | [−0.28, +0.75] |
| No MIE | [−0.06, +1.04] | [+0.04, +0.94] |

The 95% CI for WG − No MIE just barely includes zero ([−0.06, +1.04]), which is consistent with the marginally non-significant two-sided Wilcoxon $p = 0.067$ in Panel B. The 90% CI for the same comparison excludes zero ([+0.04, +0.94]), indicating that under a one-sided directional hypothesis (removing MIE files does not improve performance) the WG > No MIE direction is supported. The Cohen's $d$ on the per-question paired difference for the WG vs. No MIE comparison is 0.25.

#### Panel C. Per-condition loss summary.

| Condition | Eval. loss rate (out of 250) | Q net-loss (out of 50) | min Δ (per-q mean) | mean 5-run SD (TogoMCP) | mean worst (min-of-5) | Net-loss question IDs (Δ) |
|---|---|---|---|---|---|---|
| WG | 1.6% (4/250) | 1 | −0.20 | 0.464 | 18.02 | q021 (−0.20) |
| MIE-Instr | 4.8% (12/250) | 2 | −1.20 | 0.580 | 17.84 | q013 (−1.20), q030 (−1.00) |
| No-Instr | 6.0% (15/250) | 3 | −3.20 | 0.660 | 17.24 | q017 (−3.20), q037 (−1.00), q015 (−0.80) |
| No MIE | 4.0% (10/250) | 2 | −2.40 | 0.518 | 17.56 | q017 (−2.40), q034 (−0.20) |

*Eval. loss rate*: fraction of 250 individual question–run evaluations where the TogoMCP score fell below the matched baseline score. *Q net-loss*: number of distinct questions (out of 50) whose 5-run mean Δ was negative. *min Δ*: lowest per-question mean Δ across the 50 questions. *mean 5-run SD*: average over 50 questions of the within-question SD of the five TogoMCP total-score evaluations. *mean worst (min-of-5)*: average over 50 questions of the minimum TogoMCP total across the five evaluator runs; this is the statistic used in the worst-case paired Wilcoxon comparison reported in the main manuscript.
